# Heterologous expression of genes from a heterocystous cyanobacterial endosymbiont highlights organic carbon exchange with its diatom host

**DOI:** 10.1101/2023.01.23.525141

**Authors:** Mercedes Nieves-Morión, Sergio Camargo, Sepehr Bardi, María Teresa Ruiz, Enrique Flores, Rachel A. Foster

**Affiliations:** Department of Ecology, Environment and Plant Sciences, Stockholm University, SE-106 91 Stockholm, Sweden; Instituto de Bioquímica Vegetal y Fotosíntesis, CSIC and Universidad de Sevilla, Américo Vespucio 49, E-41092 Seville, Spain

**Keywords:** Carbon metabolism, cyanobacteria, glutamate, *Hemiaulus hauckii*, invertase, polyamines, *Richelia euintracellularis*, symbiosis

## Abstract

A few genera of diatoms are widespread and thrive in low nutrient waters of the open ocean due to their close association with N_2_-fixing, filamentous heterocyst-forming cyanobacteria. In one of these symbioses, the symbiont, *Richelia euintracellularis*, has penetrated the cell envelope of the host, *Hemiaulus hauckii*, and lives inside the host cytoplasm. How the partners interact, including how the symbiont sustains high rates of N_2_ fixation is unstudied. Since *R. euintracellularis* has evaded isolation, heterologous expression of genes in model laboratory organisms was performed to identify the function of proteins from the endosymbiont. Gene complementation of a cyanobacterial invertase mutant and expression of the protein in *Escherichia coli* showed that *R. euintracellularis* HH01 possesses a neutral invertase that splits sucrose producing glucose and fructose. Several solute binding proteins (SBPs) of ABC transporters encoded in the genome of *R. euintracellularis* HH01 were expressed in *E. coli* and their substrates were characterized. The selected SBPs directly linked the host as the source of several substrates, e.g., sugars (sucrose, galactose), amino acids (glutamate, phenylalanine) and a polyamine (spermidine), to support the cyanobacterial symbiont. Finally, transcripts of genes encoding the invertase and SBPs were consistently detected in wild populations of *H. hauckii* collected from multiple stations and depths in the western tropical North Atlantic. Our results support the idea that the diatom host provides the endosymbiotic cyanobacterium with organic carbon to fuel N_2_ fixation. This knowledge is key to understand the physiology of the globally significant *H. hauckii-R. euintracellularis* symbiosis.

**SIGNIFICANCE:** Diatom diazotroph associations (DDAs) between diatoms and N_2_-fixing bacteria (diazotrophs) have a relevant impact on N_2_ fixation-based production, but the mechanisms underlying their integrated N_2_ and CO_2_ fixation remain unstudied. In the association between the diatom *Hemiaulus hauckii* (host) and the N_2_-fixing, heterocyst-forming cyanobacterium *Richelia euintracellularis* (endosymbiont), the cyanobacterium is uncultivable. Here we used heterologous expression of genes from the endosymbiont to identify the function of proteins involved in the utilization of organic carbon from the host. The importance of these proteins was also confirmed by estimating gene expression in environmental samples. Our results show that the metabolisms of the symbiotic partners are integrated allowing the host to sustain the physiology of the endosymbiont for an important ecological role.

## INTRODUCTION

Diatom diazotroph associations (DDAs) are stable mutualistic partnerships between a few genera of diatoms and N_2_-fixing bacteria (diazotrophs). Diatoms are diverse photosynthetic unicellular microalgae that typically thrive in coastal zones with high nutrients and contribute approximately 20% of the Earth’s photosynthesis (1). The diatoms that form symbiosis with N_2_-fixing bacteria reside in regions where nutrients are sparse, and thus the symbiont provides the diatoms a new niche. Moreover, large seasonal blooms of DDAs are observed worldwide and have a direct impact on N_2_-fixation based production and carbon sequestration to the sea floor (2–4). Despite their broad distributions and significant biogeochemical impact, the molecular mechanisms regulating their integrated N_2_ and CO_2_ fixation remain unstudied.

In several open ocean DDAs, the symbionts are different species of *Richelia*, which are filamentous, terminal heterocyst-forming cyanobacteria (5). The relationships between *Richelia* and their respective diatom hosts are highly specific (6–7), and the symbiont genome size and content depend on the symbiont cellular location (8). Of interest here is *Richelia euintracellularis*, which has penetrated both components of the cell envelope (outer silicified cell wall and internal cell membrane) of its host diatom (*Hemiaulus hauckii*) (9–10) and resides in the cytoplasm of the diatom. The endosymbiont *R. euintracellularis* strain HH01 (hereafter ReuHH01; recently renamed from RintHH01; 5) possesses the smallest of all available *Richelia* genomes (3.2 Mbp), reducing in particular its N assimilatory pathways to only N_2_ fixation, and additionally lacks several central metabolic enzymes, such as glutamine:2-oxoglutarate amidotransferase (GOGAT) or the proteins for sucrose biosynthesis, and has limited transporters compared to free-living cyanobacteria (8, 11). Consistently, ReuHH01 appears unable to grow freely outside its symbiotic host (12). Genome reduction is the norm in obligate endosymbionts and suggests a high host dependency.

Important aspects of symbioses are partner function and substrate exchanges. In DDAs, the symbiont function, N_2_ fixation, is obvious and shown on the cellular level (13). However, recent evidence suggests that the diatom host provides ReuHH01 with reduced carbon (14). A closer inspection of the ReuHH01 draft genome highlights a number of potential metabolic linkages between the partners (11). For example, ReuHH01 has retained a number of ABC-type uptake transporters necessary for assimilation of sugars and amino acids. ABC uptake transporters are typically composed of a periplasmic solute-binding protein (SBP), two similar transmembrane proteins or domains (TMD) and two nucleotide-binding proteins or domains (NBP) (15). In ABC uptake transporters, the periplasmic SBP defines well the substrate specificity and is key to determine the affinity of the system for its substrate. ReuHH01 has retained homologues of the ABC transporter Gls, which functions to take up glucosides (e.g., in *Anabaena* sp. strain PCC 7120; 16), and of both N-I and N-II ABC transporters, which are responsible for amino acid uptake (11). An invertase (enzymes that irreversibly hydrolyze sucrose into glucose and fructose) is also predicted to be encoded in the ReuHH01 genome. These genomic observations suggest that ReuHH01 has retained transporters and enzymes to assimilate organic carbon received from its host.

Of additional relevance is the presence of a fully functional urea cycle in diatoms (17). Unlike the urea cycle in metazoans, the diatom urea cycle functions as an anabolic pathway producing a number of metabolic intermediates (18), some of which are necessary for both symbiotic partners. For example, polyamines such as spermidine and putrescine are derived from the diatom urea cycle. Polyamines are required by both partners: diatoms use polyamines to build their silicious cell walls and cyanobacteria require polyamines for heterocyst differentiation (19–20). Unexpectedly, the ReuHH01 genome lacks genes for a complete polyamine biosynthesis pathway (11). However, ReuHH01 has retained a possible ABC transporter homologue for polyamine import (PotADB) (11). Thus, the diatom host potentially plays a larger role in the symbiont’s overall function, including the provision of organic carbon.

Since ReuHH01 has evaded isolation, we used the heterologous expression of ReuHH01 genes of interest in laboratory amenable organisms, including *Escherichia coli* and another heterocyst-forming cyanobacterium, *Anabaena* sp. strain PCC 7120 (hereafter *Anabaena*), in order to identify gene function. Here, we studied properties of several SBPs from ABC transporters of ReuHH01 putatively involved in uptake of organic carbon and additionally properties of an invertase from this organism. We then showed gene expression for the genes that encode some components of these ABC transporters and invertase in wild symbiotic populations. Knowledge of these proteins is key to understand the physiology of the globally significant *H. hauckii-R. euintracellularis* symbiosis.

## RESULTS

### ReuHH01 possesses an SBP with high affinity for sucrose

The ABC transporter Gls of *Anabaena* mediates the uptake of glucosides and some monosaccharides such as glucose and fructose (16). Genes encoding homologues to *Anabaena* Gls transporter components could be identified in the ReuHH01 genome (11; summarized here in Table S1 in Supporting Information; RintHH01 was recently renamed ReuHH01, but gene/protein homologues hereafter retain original epithet to facilitate their identification in the databases; 5). The predicted periplasmic SBP, RintHH_17430, shows limited similarity to *Anabaena* GlsR, with which it shares 23% identity (87/376 amino acid residues). We focused on the analysis of RintHH_17430 to address the substrate specificity of the transporter. RintHH_17430 (gene ID, 2580948285) was chemically synthesized and cloned in *E. coli* expression vector pET28b(+) (Fig. S1). Because the RintHH_17430 protein has a signal peptidase II-type signal peptide (Fig. S2A), which determines processing of the protein with addition of a lipid to a cysteine (Cys) residue (21), we cloned the gene to position the mature protein with the N-terminal Cys residue fused to the Strep tag II, thus avoiding anchoring to the cytoplasmic membrane and facilitating the purification of the protein (Fig. S1). The mature RintHH_17430 protein has 12 tryptophan (Trp) residues (Fig. S2B), which permits to test binding of putative substrates by fluorescence quenching (22).

Purified RintHH_17430 protein excited by light at 280-nm showed strong fluorescence from 300 to 400 nm, which corresponds to Trp fluorescence (Fig. 1A). This fluorescence was efficiently quenched (about 22.5% quenching) by sucrose (glucose 1α➔2 fructose) at 1 μM. Increasing sucrose concentration to 1 mM increased quenching only to about 35% (Fig. 1A), indicating a high affinity of the protein for sucrose. Other sugars, glucose, fructose, maltose (glucose 1α➔4 glucose), galactose and, for comparison, an amino acid, glutamate, were also tested. In general, fluorescence quenching was observed, but it was lower than the quenching for sucrose at the lower concentrations (1 and 10 μM; Fig. 1B). The fact that 1 μM glutamate produced quenching similar to sugars other than sucrose suggests low-specificity binding for several compounds. However, 10 μM to 1 mM galactose produced higher quenching than glucose, fructose, maltose, and glutamate; this quenching was also close to the quenching by sucrose at the highest concentration tested (1 mM). The extent of RintHH_17430 sucrose-induced fluorescence quenching is similar or somewhat stronger than that described for other periplasmic binding proteins and their substrates (see, e.g., 22-24) These results suggest that the genome of ReuHH01 encodes an ABC transporter with a periplasmic SBP (GlsR) that can bind sucrose with high affinity and other sugars such as galactose with lower affinity.

**Fig. 1.**
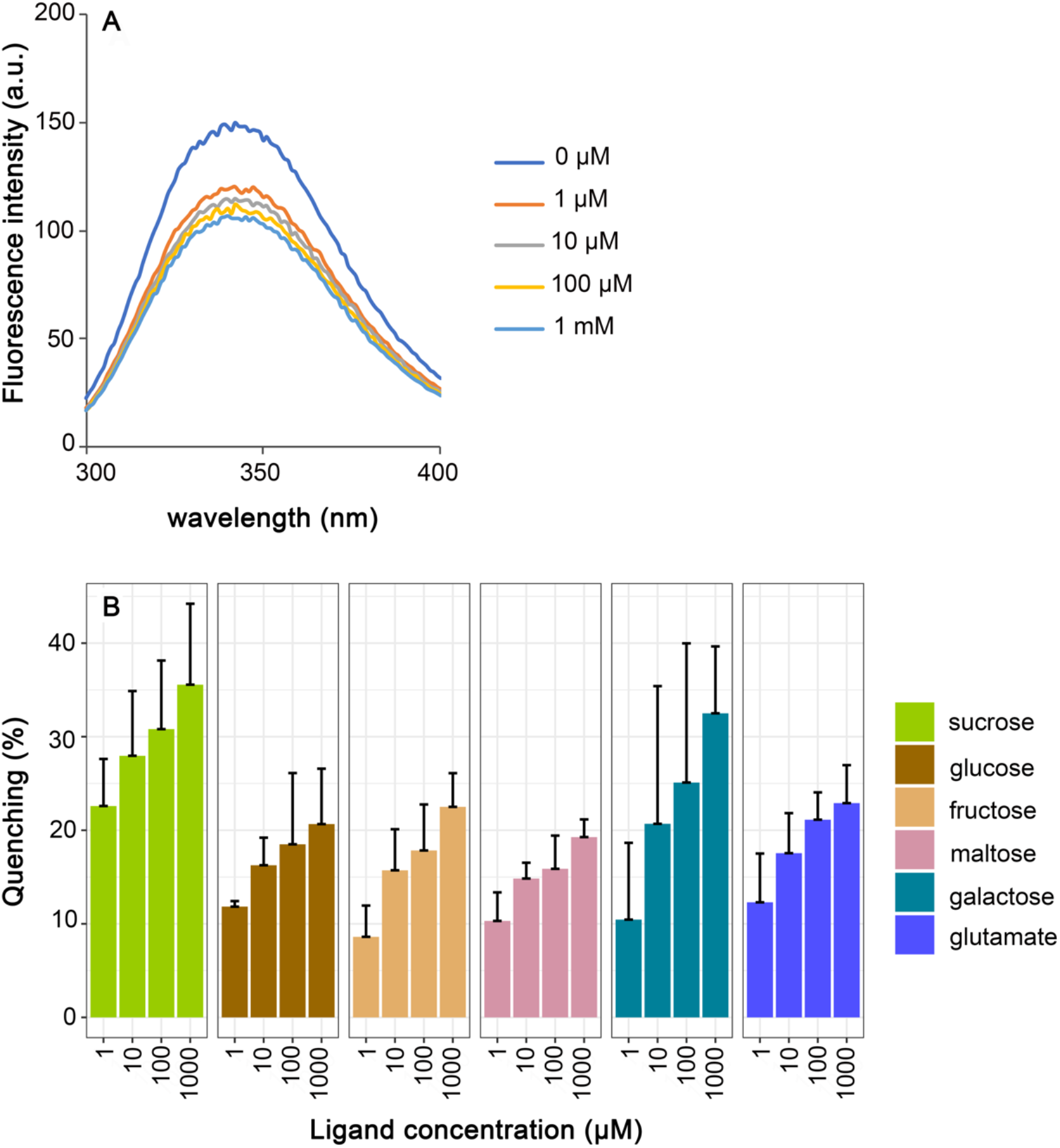
Substrate tests of the periplasmic solute-binding protein RintHH_17430 from *Richelia euintracellularis*. (A) Changes in intrinsic tryptophan fluorescence of RintHH_17430 upon the addition of sucrose at the indicated concentrations. RintHH_17430 (0.02 mM) was excited at 280 nm and the emission spectrum from 300 to 400 nm was recorded. A representative example is shown. (B) Quenching of RintHH_17430 fluorescence in response to different compounds. The fluorescence emission of RintHH_17430 was followed at the maximal emission wavelength (around 340 nm) in the absence of added sugar (or glutamate) or after the addition of 1, 10, 100 and 1000 mM of sucrose, glucose, fructose, maltose, glutamate or galactose. Data are the mean and standard deviation (at least three replicates) of the quenching induced by a compound at the indicated concentration (i.e., % decrease of the fluorescence observed in the absence of any metabolite).

### ReuHH01 possesses a neutral invertase for splitting sucrose

ORF RintHH_3860 of ReuHH01 (gene ID, 2580946898) encodes a protein that shows homology to invertases: 74% identity (347/467 amino acid residues) to *Anabaena* Alr0819 (InvB); and 60% identity (272/456 amino acid residues) to *Anabaena* Alr1521 (InvA) (Fig. S3A). Because the *Anabaena invB* mutants are unable to grow fixing N_2_ under oxic conditions (25, 26), we pursued complementation of an *Anabaena invB* mutant with RintHH_3860 to test the possible role of this ReuHH01 gene product as an invertase. For this, the RintHH_3860 gene was chemically synthesized together with the *Anabaena invB* gene promoter (Fig. S3B) and cloned in a plasmid, and the *invB* mutant described previously by López-Igual et al. (25) was re-constructed (see Fig. S4). The RintHH_3860 gene construct was transferred by conjugation to the *Anabaena invB* mutant producing a strain in which RintHH_3860 substituted for *invB* and was expressed from the *invB* promoter (Fig. S4). Growth tests in solid and liquid media confirmed that the *Anabaena invB* mutant could not grow diazotrophically (incubation in BG11_0_ medium) whereas it grew well in nitrate-containing (BG11) medium; in contrast, the *invB* mutant complemented with RintHH_3860 could grow fixing N_2_ as well as in nitrate-containing medium (Fig. 2A and B). Determination of the growth rate constants under diazotrophic conditions showed a limited growth rate of the *Anabaena invB* mutant (growth rate constant, 0.089 d^-1^) and a growth rate of the RintHH_3860-complemented *Anabaena* strain (0.342 d^-1^) similar to the growth rate of wild-type *Anabaena* (0.366 d^-1^) (Fig. 2C). These results identify RintHH_3860 as a functional invertase.

**Fig. 2.**
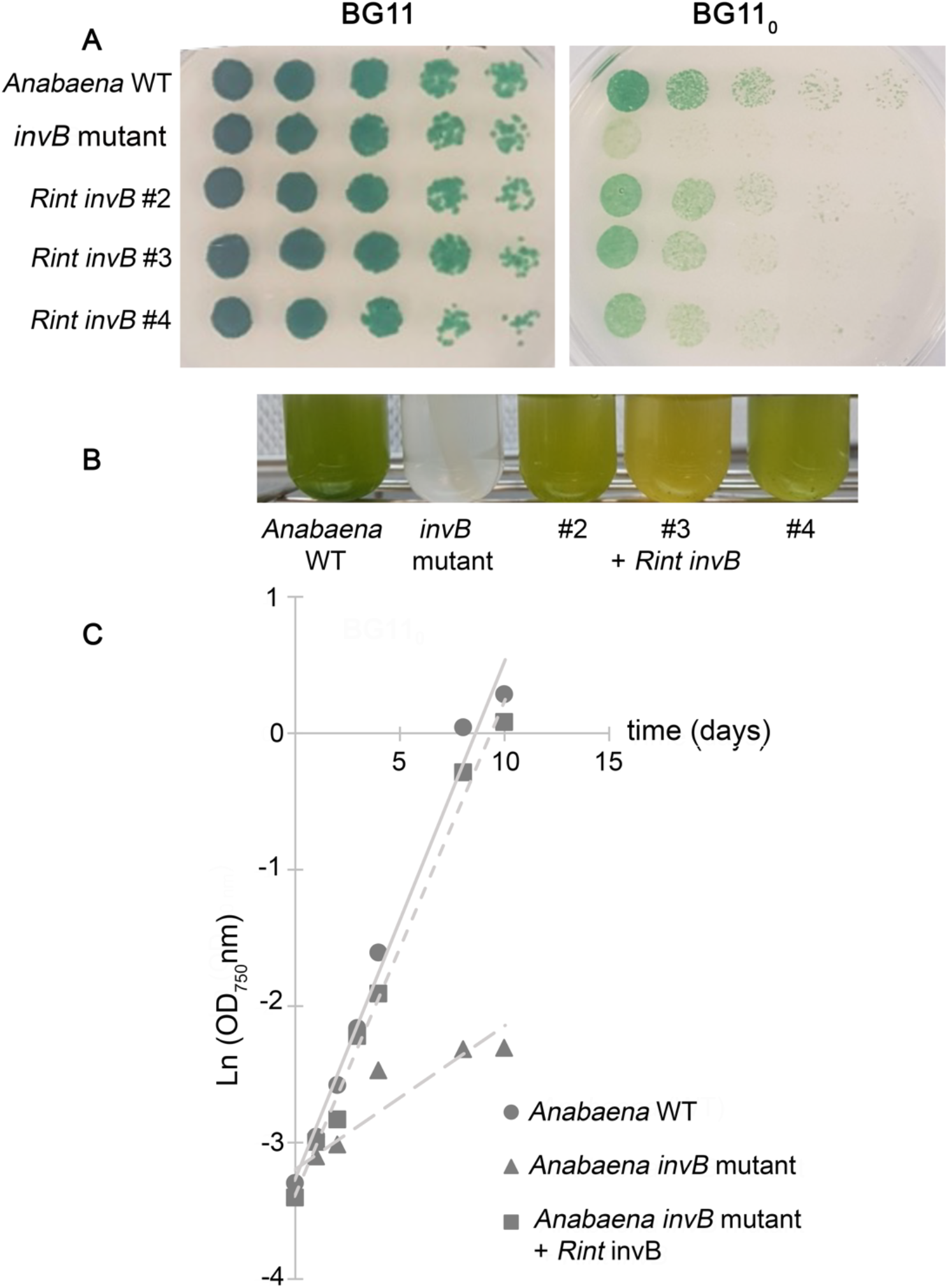
Growth tests of wild-type *Anabaena*, the *Anabaena invB* mutant and the *invB* mutant complemented with the *Richelia euintracellularis* gene RintHH_3860. **(A)** Growth test on solid medium of wild-type (WT) *Anabaena*, the *invB* mutant and the complemented strain (3 clones). Filaments grown in BG11 medium (in the presence of antibiotics for the mutants) were resuspended in BG11_0_ medium, dilutions were prepared, and a 10 μl portion of each dilution (from left to right, 1, 0.5, 0.25, 0.125, and 0.0625 μg Chl ml^-1^) was spotted on BG11 (nitrate) or BG11_0_ (N_2_) medium. The plates were incubated under culture conditions, and photographs were taken after 10 days of incubation. **(B)** Liquid growth of WT *Anabaena*, the *invB* mutant and the complemented strain in BG11_0_ medium supplemented with 1% CO_2_. Inoculum: 0.2 μg Chl ml^-1^; incubation time, 10 days. **(C)** Determination of the growth rate constant for WT *Anabaena*, the *invB* mutant and the complemented strain (one clone shown as example) in BG11_0_ medium supplemented with 1% CO_2_. Inoculum: 0.2 μg Chl ml^-1^. Growth rate constants (mean of two experiments): WT, 0.366 day^-1^; *invB* mutant, 0.089 day^-1^; complemented strain, 0.342 day^-1^.

To study the properties of the RintHH_3860 protein, the gene was cloned in an *E. coli* expression vector, pET28b(+) (Fig. S5). Because purification of the protein was unsuccessful, we studied the invertase activity in cell-free extracts of *E. coli* [pET28b(+)::RintHH_3860] compared with the activity in extracts of an isogenic strain lacking RintHH_3860. Cell-free extracts of *E. coli* bearing RintHH_3860 supplemented with sucrose showed equimolar production of glucose and fructose, indicating that both monosaccharides resulted from sucrose hydrolysis, whereas no production was observed in the extracts of the control strain (Fig. 3A). As determined in five independent assays, production was linear with time for at least 60 min. To characterize the functional group of invertases to which RintHH_3860 belongs, the pH dependency of the reaction was determined, showing an optimal pH of 7 to 7.5 (Fig. 3B). Finally, the *K*_m_ of RintHH_3860 for sucrose was determined as 25.7 mM (Fig. 3C), which is comparable to the *K*_m_ previously reported for other cyanobacterial invertases, 15 to 36 mM (27–28). It is, however, somewhat higher than the *K*_m_ reported for *Anabaena* InvB, 9 to 15 mM (27; 29). Thus, RintHH_3860 is a neutral invertase and, consistent with its homology to *Anabaena* InvB, we term it InvB.

**Fig. 3.**
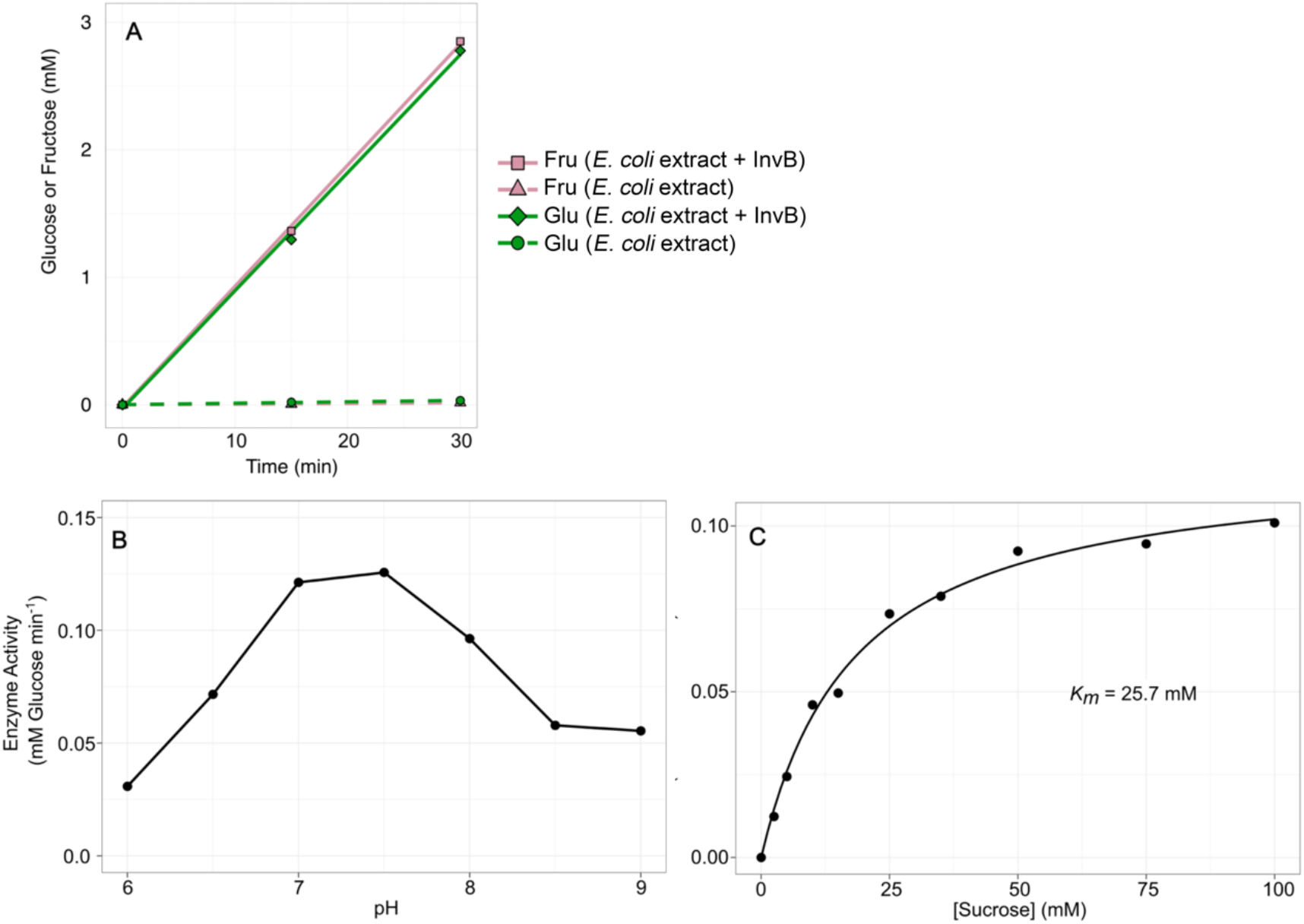
Characterization of RintHH_3860 from *Richelia euintracellularis* as a neutral invertase. **(A)** Enzyme activity in crude extracts from *E. coli* and *E. coli* + InvB. Substrate (sucrose) concentration, 100 mM; pH 7. Cell extracts, 1.5 (*E*. coli) or 1.3 (*E. coli* + InvB) μg of protein ml^-1^. The production of glucose and fructose was determined and their concentration in the assay along time is shown. **(B)** Enzyme activity of invertase at different pH values. Substrate (sucrose) concentration, 100 mM; cell extracts, 1.5 μg of protein ml^-1^. Rate of glucose production is shown. Data represent the means of two independent assays. The results show that InvB is a neutral invertase. **(C)** Determination of the *K*_m_ for sucrose of InvB. Rate of glucose production at different initial concentrations of sucrose is shown; data represent the mean of two independent assays. Cell extracts, 1.5 μg of protein ml^-1^; pH, 7; rates calculated from 30- and 60-min incubation times, which showed lineal production of glucose.

### ReuHH01 possesses SBPs with variable affinities for amino acids and polyamines

The genome of ReuHH01 contains genes putatively encoding the components of ABC uptake transporters that are homologs, respectively, to the *Anabaena* N-I transporter for neutral/hydrophobic amino acids and N-II transporter for acidic and neutral amino acids (11; see Table S1). The ReuHH01 genome also putatively encodes the components of an ABC uptake transporter homolog to the *Anabaena* Pot transporter for polyamines, although the putative substrate binding protein shows relatively low similarity (11; see Table S1). We then addressed the study of the periplasmic substrate-binding proteins of these transporters. The three *R. euintracellularis* genes, RintHH_11820 (gene ID 2580947711), RintHH_12770 (gene ID 2580947807), and RintHH_7180 (gene ID 2580947237), encode proteins that contain a signal peptidase II-type of signal peptide and Trp residues (Fig. S6). The genes were chemically synthesized and cloned in a plasmid, and each of the predicted mature proteins (lacking the signal peptide) was produced in *E. coli* with an N-terminal His tag and purified (Figs. S7, S8, S9). When excited with light of 280 nm, the proteins showed Trp fluorescence (Figs. S7, S8, S9), which permitted to test substrate-induced fluorescence quenching as described earlier for the ReuHH01 GlsR protein.

RintHH_11820 and RintHH_12770 are homologs of the *Anabaena* NatB and NatF proteins (SBP of the N-I transporter and SBP of the N-II transporter, respectively). Fluorescence quenching of RintHH_11820 was tested by addition of the following compounds: L-Pro, L-Ala, L-Phe, Gly, L-Leu, L-Asp, L-Arg and glucose. Added at ca. 116 μM, substantial quenching was produced by the neutral amino acids (Pro, Ala, Phe, Gly, Leu) but much less quenching was produced by the sugar or by the acidic (Asp) or basic (Arg) amino acids, and ca. 17 μM Phe was especially effective denoting high affinity (Fig. 4A). We conclude that RintHH_11820 (NatB) is the binding protein of an ABC transporter similar to the *Anabaena* transporter N-I for neutral/hydrophobic amino acids (30). However, other amino acids remain to be tested for binding by RintHH_11820; those presented here were expected for the diatom cytoplasm. Similarly, fluorescence quenching of RintHH_12770 was tested by addition of the following compounds: L-Glu, L-Asp, L-Leu, and glucose. At ca. 7 μM concentration, substantial quenching was only seen with Glu; however, at 120 μM and higher concentrations, also Asp showed substantial quenching (Fig. 4B). Only low quenching was observed with Leu or glucose. These results identify RintHH_12770 as a high-affinity glutamate binding protein, NatF, and suggest that the corresponding ABC transporter of ReuHH01 is similar but not identical to the *Anabaena* N-II transporter, which is more important in aspartate transport than in glutamate transport (31).

**Fig. 4.**
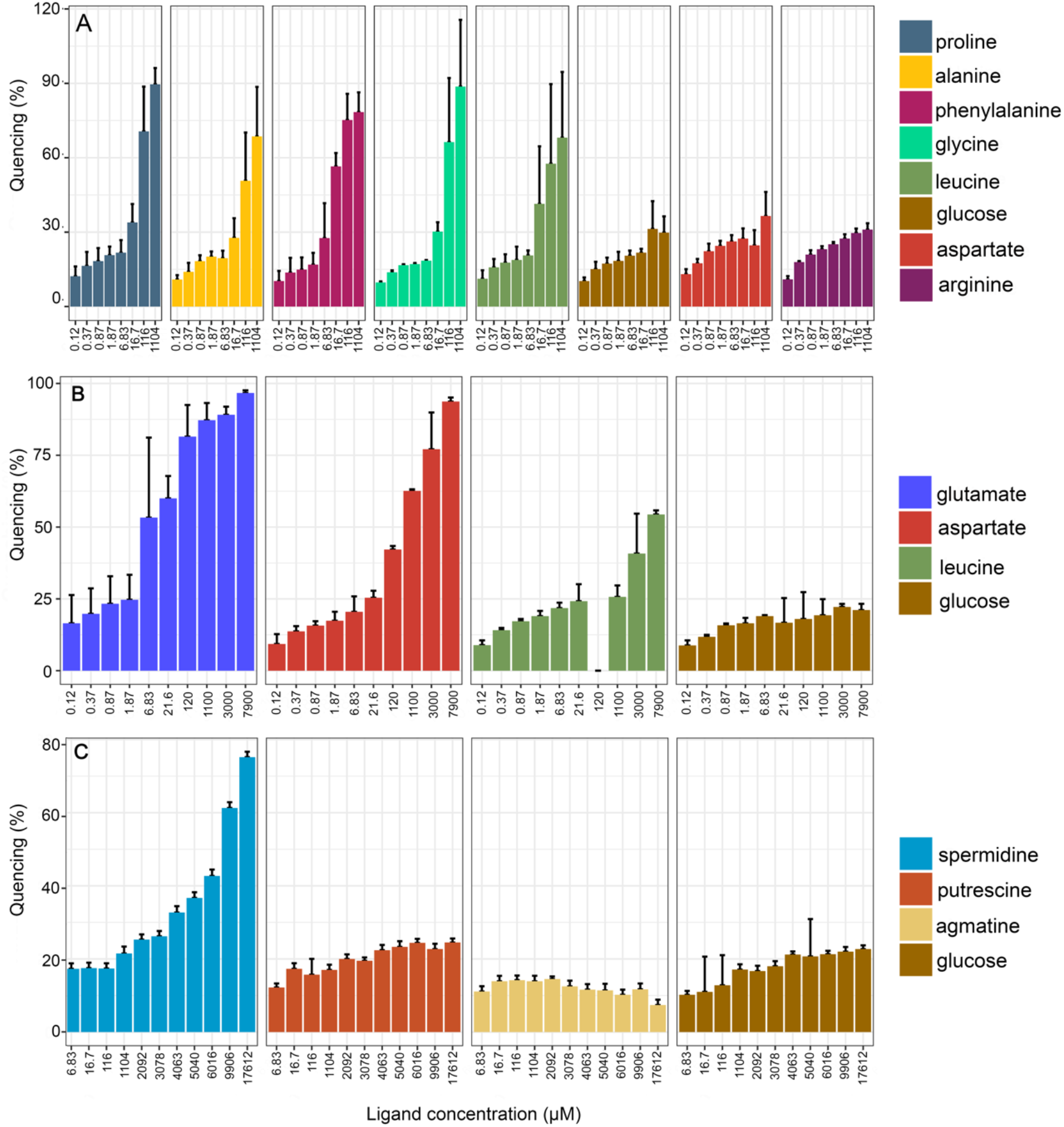
Substrate tests of different periplasmic solute-binding proteins (SBPs) related to nitrogen metabolism from *Richelia euintracellularis*. Quenching of three SBPs fluorescence in response to different nitrogen compounds and glucose as a negative control. The fluorescence emission of the three SBPs was followed at the maximal emission wavelength (around 340 nm) in the absence of substrate or after the addition of a wide range of substrate concentrations. Plotted data are the mean and standard deviation (between two and four replicates) of the quenching induced by a substrate at the indicated concentration. **(A)** Quenching of RintHH_11820 fluorescence in response to different amino acids and glucose. **(B)** Quenching of RintHH_12770 fluorescence in response to different amino acids and glucose (leucine at 120 μM was not tested). **(C)** Quenching of RintHH_7180 fluorescence in response to different polyamines and glucose.

RintHH_7180 shows homology to *Anabaena* PotD (SBP of the Pot transporter; 20), albeit similarity is relatively low (Table S1). Fluorescence quenching was tested by addition of the following compounds: agmatine, putrescine, spermidine, and glucose. None of these substrates produced significant quenching when added at 1.1 mM concentration or lower. However, when spermidine was added at concentrations greater than 3 mM, substantial quenching was produced, whereas the three other compounds produced only a basal effect (Fig. 4C). These results show that RintHH_7180 (PotD) is a low-affinity binding protein for spermidine, although the possibility remains that it has higher affinity for a non-tested polyamine.

### Environmental expression of *R. euintracellularis glsR, invB, natF, and potD*

To estimate the possible *in situ* activity of the proteins studied above in the *H. hauckii-R. euintracellularis* symbiosis in a natural environment, we quantified gene transcripts from select locations in the Western Tropical North Atlantic (WTNA). Specific oligonucleotides and assays were designed to detect gene transcripts similar to ReuHH01 *invB, glsR, natF* and *potD;* and *secA*, encoding protein translocase subunit SecA, was used for normalization (Table S2). Importantly, the sampling occurred during a high-density bloom of *H. hauckii-R. euintracellularis* over a 3-week period (32). The selected stations were chosen based on ship-board observations of high densities of *H. hauckii-R. euintracellularis*, and they represented geographically and temporally distinct locations. For example, station 2 is in close proximity (24 km) to stations 23 and 25, but was sampled 3 weeks earlier, while stations 2 and 5 were sampled 3 days apart but more distantly located (557 km) (Fig 5A).

**Fig. 5.**
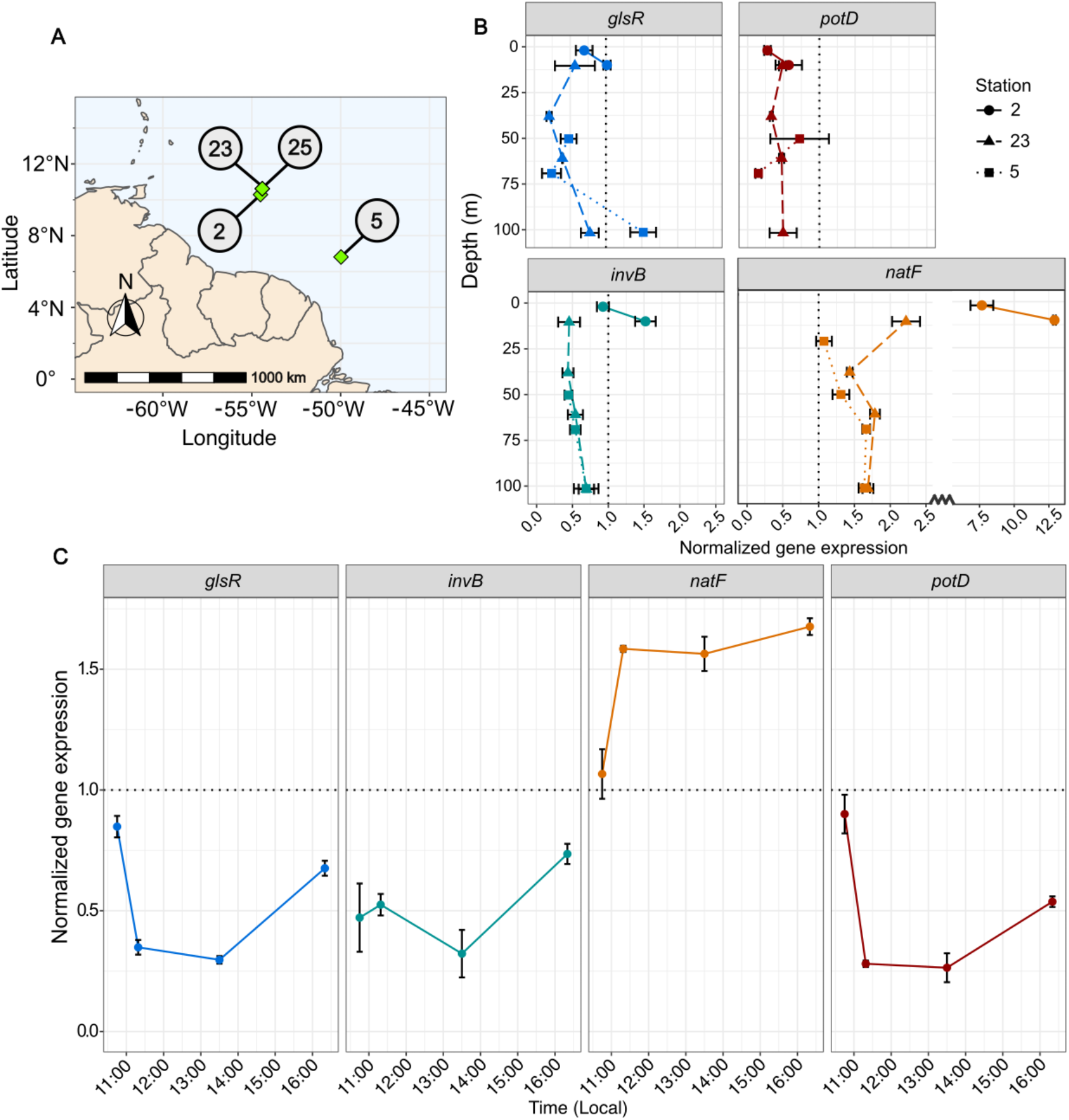
Environmental expression of genes related to nitrogen and carbon metabolism in *Richelia euintracellularis*. Gene expression for *glsR, invB, natF* and *potD* was quantified in samples from the WTNA with RT-qPCR assays normalized to the *secA* gene. **(A)** Map of the sampling locations. **(B)** Depth profiles of normalized gene expression for the respective genes at select stations (2, 5, 23). The samples were taken in the early light period between 08:00-11:30 (local time). **(C)** Time series of expression of gene targets during the light period in samples collected at station 25 (Depth = 30 m). All error bars represent the standard error of the mean of triplicate reactions in the qPCR assays. Samples for which only one or two reactions showed detection were not included.

Transcripts of all target genes were detected at different depths in the water column and at all stations (Fig. 5B; Table S3). The transcripts of *natF* were detected at particularly significant levels compared to the gene chosen as reference, *secA* (Fig. 5B; Table S3), and expression remained high throughout the daylight at station 25 (Fig 5C; Table S3). Notably, *glsR* and *invB* transcripts were detected at similar levels throughout the light period (Fig. 5C; Table S3), consistent with a related function of their protein products. Expression of *potD* was similarly low as *glsR* and *invB* but consistently detected at multiple depths, and at highest levels earlier in the light period (Fig. 5B, 5C; Table S3).

## DISCUSSION

*Richelia euintracellularis* represents one of the most widespread heterocyst-forming cyanobacteria in the world’s oceans, and it does so thriving inside the cytoplasm of its diatom host *Hemiaulus hauckii* (10). *R. euintracellularis* can fix atmospheric N_2_ at levels >100-fold higher than those exhibited by free-living heterocyst-forming cyanobacteria (13). A filament of *R. euintracellularis* represents about 1 % of the volume of a host diatom, and two *R. euintracellularis* filaments are routinely observed per *H. hauckii* host (10). Evidence suggesting that the host diatom transfers organic carbon to the endosymbiont has recently been obtained (14), and transfer is also predicted based on a hypothetical flux model (33). This raises a scenario in which the diatom can help to support the high N_2_ fixation rates of the cyanobacterium by transfer of reduced (organic) carbon compounds, which are needed to fuel N_2_ fixation in the heterocysts (34).

The genome of ReuHH01 encodes a number of possible membrane transporters that may be involved in uptake of organic compounds from the diatom host (11). The study of the substrates of those transporters, or, generally, of proteins from *R. euintracellularis*, is however hampered by the inability of this organism to grow in the laboratory, perhaps as a consequence of its reduced genome (8). We therefore approached the study of some genes encoding proteins that might be relevant in the organic nutrition of ReuHH01 by their expression in heterologous systems. Studying periplasmic solute binding proteins from ReuHH01, we have provided evidence for the presence in the endosymbiont of ABC transporters that can take up sucrose, glutamate and neutral/hydrophobic amino acids, respectively, with high-affinity. Additionally, there is a low-affinity but high-specificity transporter for spermidine. The genes which encode these transporters are also expressed by wild populations in the WTNA, demonstrating *in-situ* activity.

ReuHH01 appears to have an ABC uptake transporter similar to the *Anabaena* Gls system for glucosides and some monosaccharides (11). Its periplasmic binding protein, RintHH_17430 (GlsR), binds with high affinity sucrose and, with lower affinity, galactose at significant levels, and glucose, fructose and maltose at lower levels. At least glucose and galactose have been described to be found at substantial levels in diatoms (35), suggesting that these monosaccharides could be incorporated into the endosymbiont from the host. Glucose can be used in the heterocyst to fuel N_2_ fixation (36), and ReuHH01 has at least one enzyme of galactose catabolism, UDP-glucose 4-epimerase (GalE) (RintHH_13420). Dissolved neutral sugars comprise a reactive component of the dissolved organic carbon pool (DOC) in seawater, and thereby make concentration measurements challenging. However, in samples collected from the oligotrophic Sargasso Sea, an area where DDAs are known to occur, glucose, galactose, fucose, and mannose + xylose, are the main sugar constituents, and in the surface, dissolved combined sugar concentrations varied between 0.5-0.8 μM (37–38). Some diatoms have sugar transporters, raising the possibility of uptake from the environment (17). Thus, it is possible that a dissolved pool of sugars is available to the host diatom, in addition to the sugars produced from photosynthesis.

On the other hand, sucrose is generally considered not to be synthesized by diatoms. Because ReuHH01 also seems unable to synthesize sucrose (11), the question arises of the source of sucrose for invertase, which is encoded in the ReuHH01 genome and expressed by the wild populations as shown in this work. Interestingly, the model diatom *Phaeodactylum tricornutum* has one SWEET protein (TCDB 2.A.123.1.15), a type of transporter involved in the uptake of sugars including sucrose (39). Sucrose cleaved by invertase is involved in N_2_ fixation (25–26, 29, 40) producing glucose and fructose that support heterocyst metabolism (35). Further work will be necessary to clarify sucrose metabolism in *R. euintracellularis* and its relation to the metabolism of *H. hauckii*.

ReuHH01 has been shown to contain glutamine synthetase (GS) but to lack glutamine:2-oxoglutarate amidotransferase (GOGAT), raising the question of the origin of glutamate that is used to incorporate ammonium resulting from N_2_ fixation (8). It was proposed that glutamate dehydrogenase (normally a glutamate catabolism enzyme; 41) encoded in the ReuHH01 genome could be involved in ammonium assimilation (8). However, the presence of a high-affinity glutamate transporter in the endosymbiont, as shown in this work, and of high concentrations of glutamate in the cytoplasm of most organisms including diatoms (35), raises the possibility that ReuHH01 obtains glutamate from its host. Glutamate could then be used by GS to incorporate ammonium and in many other metabolic reactions, including catabolism by glutamate dehydrogenase, in the endosymbiont. Moreover, the *in-situ* transcriptional activity for *natF* showed the highest expression of all genes analyzed, and coincides with the time of day when N_2_ fixation is expected to be highest (12).

*R. euintracellularis* HH01 also encodes an ABC uptake transporter similar to the N-I system of *Anabaena*, which transports neutral/hydrophobic amino acids (30). Here, we have shown that the periplasmic binding protein of this transporter binds with high affinity phenylalanine, and with somewhat lower affinity alanine, glycine, leucine and proline. All these amino acids are found at substantial levels in diatoms (35), suggesting that the endosymbiont can obtain a supply of some amino acids from the host. In *Anabaena*, alanine is used by an alanine dehydrogenase that is expressed in the heterocysts supporting diazotrophic growth (42). The ReuHH01 genome contains a gene (RintHH_16150) that encodes a protein with 80 % identity to alanine dehydrogenase of *Anabaena*, supporting a possible role of alanine taken up from the host in N_2_ fixation by ReuHH01. Proline is also of interest because it is synthesized as an osmolyte from the urea cycle that is active in diatoms (18), and some of the proline accumulated under osmotic stress could therefore be incorporated by the neutral/hydrophobic amino acid ABC transporter into the endosymbiont to support protein synthesis.

In diatoms, the urea cycle is also a source of polyamines (18), which are needed, as long-chain polyamines, to build the frustule (43). As precursors of the long-chain polyamines, the diatoms accumulate polyamines such as spermidine (45). The heterocyst-forming cyanobacteria require polyamines to complete heterocyst differentiation (20), and homospermidine is the polyamine characteristic of heterocyst-forming cyanobacteria (45). However, ReuHH01 lacks a complete polyamine biosynthesis pathway (11). The presence of an ABC transporter whose periplasmic binding protein can bind spermidine, albeit with relatively low affinity, suggests that the endosymbiont can obtain polyamines from the host. To this end, we also detected expression of *potD* in the wild populations. Whether *R. euintracellularis* can use spermidine as a substitute for homospermidine is unknown, but we note that, for example, *Anabaena* seems to be able to use alternative polyamines when homospermidine synthesis has been inactivated (20).

Here, we have shown that specific genes from ReuHH01 encode proteins that can bind sugars, amino acids and a polyamine, respectively. Additionally, we have demonstrated that a ReuHH01 gene encodes a neutral invertase. Importantly, transcripts of all those genes were detected in environmental samples of a *H. hauckii-R. euintracellularis* bloom, indicating that the genes are active in producing proteins that can mediate assimilation of organic carbon from the host. Some of the metabolites taken up by *R. euintracellularis* may have specific metabolic or physiological roles, such as glutamate in ammonium assimilation, polyamines in heterocyst differentiation, and alanine or the sugars supporting N_2_ fixation. In summary, our results strongly support the idea that *R. euintracellularis* expresses transporters and enzymes that can mediate the use of organic carbon obtained from the host diatom to fuel N_2_ fixation (Fig 6). Our work presented here on membrane transport, combined with the earlier observations of an adapted cell envelope in ReuHH01 (e.g., presence of vesicles, proximity to mitochondria) (10), establishes further directions for research on these biogeochemically relevant populations.

**Fig. 6.**
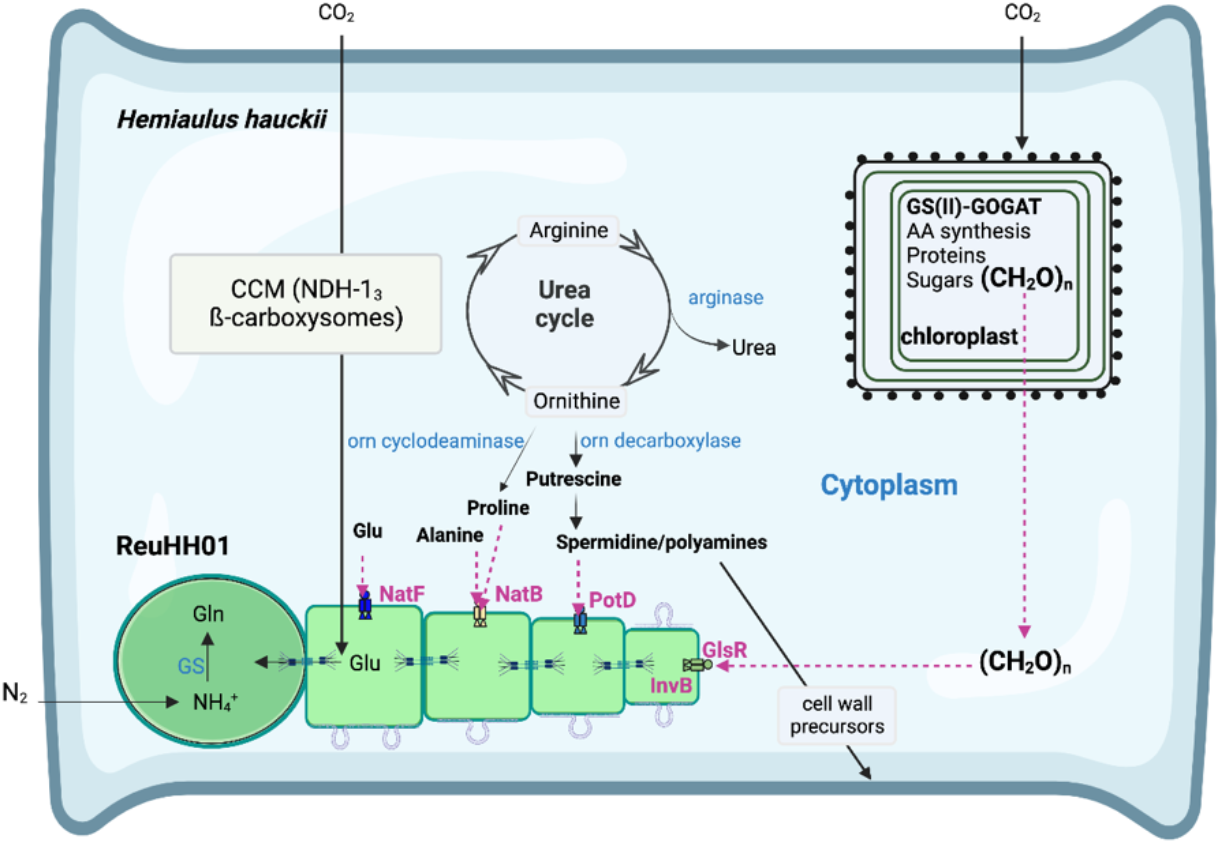
Summary of potential metabolic exchanges in the *Hemiaulus hauckii-Richelia euintracellularis* (ReuHH01) symbioses presented in this work. Both partners can fix CO_2_ photosynthetically and additionally the host appears to supply the symbiont with organic carbon to fuel its high N_2_ fixation rate in the heterocyst (left terminal cell in the ReuHH01 filament). Typically, two ReuHH01 are found per host (5), here only one is shown for simplicity. The urea cycle is known to be operative in diatoms (18). The transporters whose SBPs have been characterized in this work can be involved in uptake by ReuHH01 of amino acids such as glutamate (NatF), alanine or proline (NatB), a polyamine (spermidine; PotD) and sugars (GlsR). A neutral invertase, InvB, is also expressed in ReuHH01; here it is depicted as located in vegetative cells, but it could be specifically located in the heterocyst as in *Anabaena* (25). Metabolites are homogenized along the cyanobacterial filament by diffusion through proteinaceous septal junctions represented as septal structures (48). The endosymbiont provides the host with fixed nitrogen, but the mechanism of transfer is unknown. Also depicted are numerous membrane vesicles formed on the cell envelope of vegetative cells of ReuHH01 (here shown as vesicles in formation), which could potentially function in metabolite transfer between partners (10). AA, amino acids; CCM, carbon concentration mechanism; Glu, glutamate; Gln, glutamine; Orn, ornithine; GS, glutamine synthetase; GOGAT, glutamine:2-oxoglutarate amidotransferase. Structures not to scale.

## MATERIALS AND METHODS

### Genomic information

The draft genome sequence of *Richelia euintracellularis* HH01 is available at https://img.jgi.doe.gov/cgi-bin/m/main.cgi. It should be noted that since the release of the genome sequence, the *Richelia* species has been updated (5); the previous name for *Richelia euintracellularis* was *Richelia intracellularis*. The proteins described in this work can be downloaded from https://www.uniprot.org. Protein sequence comparisons were done by BlastP (https://blast.ncbi.nlm.nih.gov/Blast.cgi?PAGE=Proteins) and assessed at the Phyre2 portal for protein modeling, prediction and analysis (http://www.sbg.bio.ic.ac.uk/~phyre2/html/page.cgi?id=index). Amino acid composition of a protein was analyzed at https://web.expasy.org/protparam/. Signal peptides were searched at http://www.cbs.dtu.dk/services/SignalP/.

### Expression and purification of SBPs

The genes were synthesized by Integrated DNA Technologies, Inc. (IDT), and provided in a plasmid from which the selected DNA fragment was transferred by standard recombinant DNA methods to the plasmid shown in the corresponding supplementary figures (Figures S1; S4-S5; S7-S9). Oligodeoxynucleotide primers used in plasmid constructions are shown in Suppl. Table S4. Proteins were produced in *E. coli* after standard IPTG-induction and purified by affinity chromatography using StrepTrap™ HP column (prepacked with StrepTactin Sepharose) or nickel columns for proteins carrying Strep-tag II or 6xHis fusions, respectively. See Suppl. Methods for details on protein purification. Protein fluorescence was induced at 280 nm and the emission recorded between 300-400 nm in a Varian Cary Eclipse Fluorescence Spectrophotometer. Maximum fluorescence intensity was at 340 nm and used to estimate the percent (%) quenching for each ligand tested.

### Invertase analysis

The *Anabaena invB* mutant described in López-Igual et al. (25) was re-constructed and complemented with RintHH_3860 (synthetic gene provided by IDT) from ReuHH1 as described in Fig. S4. DNA constructs were transferred to *Anabaena* by triparental conjugation and *Anabaena* was grown photo-autotrophically as previously described (25). RintHH_3860 was also expressed in *E. coli* as described in Fig. S5. Invertase assays were carried out at 30 °C in 120 mM potassium-phosphate buffer of the indicated pH. Glucose and fructose produced in the reaction were detected and quantified in a Dionex ICS-5000 HPLC chromatograph equipped with CarboPac PA10 column and pre-column (46).

### Environmental sampling and gene expression by qRT-PCR

An expedition to the western tropical north Atlantic (WTNA) was conducted in May-June 2010. Four stations (2, 5, 23, 25) were selected for further analyses of gene expression based on high densities of symbiotic *H. hauckii-R. euintracellularis* from ship-board microscopy observations and previously published information (14, 32). Seawater collection from discrete depths and times used niskin bottles arranged on a conductivity depth temperature (CTD) rosette (Suppl. Table S3). Water was collected into bleach-rinsed (10 %) bottles (2.5 L) and filtered immediately onto 0.2 μm pore size filters (25-mm diameter) using a peristaltic pump. Filters were placed in sterile tubes amended with RLT buffer (Qiagen RNA easy) and stored at −80 °C until extraction. Total RNA was extracted and reverse transcribed and used in quantitative polymerase chain (RT-qPCR) reactions using previously described methods, commercially available kits (Qiagen RNA easy extraction, Superscript™ First Strand Synthesis System) and TaqMAN chemistry (Applied Biosystems) (47). The specificity of the RT-qPCR assays were evaluated *in silico* with BLASTn (Suppl. Methods). The results indicated the assays are specific to *R. euintracellularis* HH01 (Fig. S10). Further details, including the designing of the new oligonucleotides for evaluating the gene expression of *glsR, invB, natF, potD, and secA* are described in the Suppl. Methods.

## ACKNOWLEDGMENTS

Work in Stockholm and Seville was supported by grant no. 2018-04161 from The Swedish Research Council (Vetenskapsrådet) to RAF and EF. RAF is additionally funded by the Knut and Alice Wallenberg Foundation.

## AUTHOR CONTRIBUTIONS

MN-M designed and performed the research and analyzed data; SC designed and performed protein work and analyzed data; SB designed oligonucleotides and performed qRT-PCR assays and analyzed data; MTR contributed analytical tools; RAF collected samples and performed RNA extractions; EF and RAF conceived the study, supervised research and co-wrote the paper with input from all co-authors.

