## Supplemental Figures for "Heterologous expression of genes from a heterocystous cyanobacterial endosymbiont highlights organic carbon exchange with its diatom host"

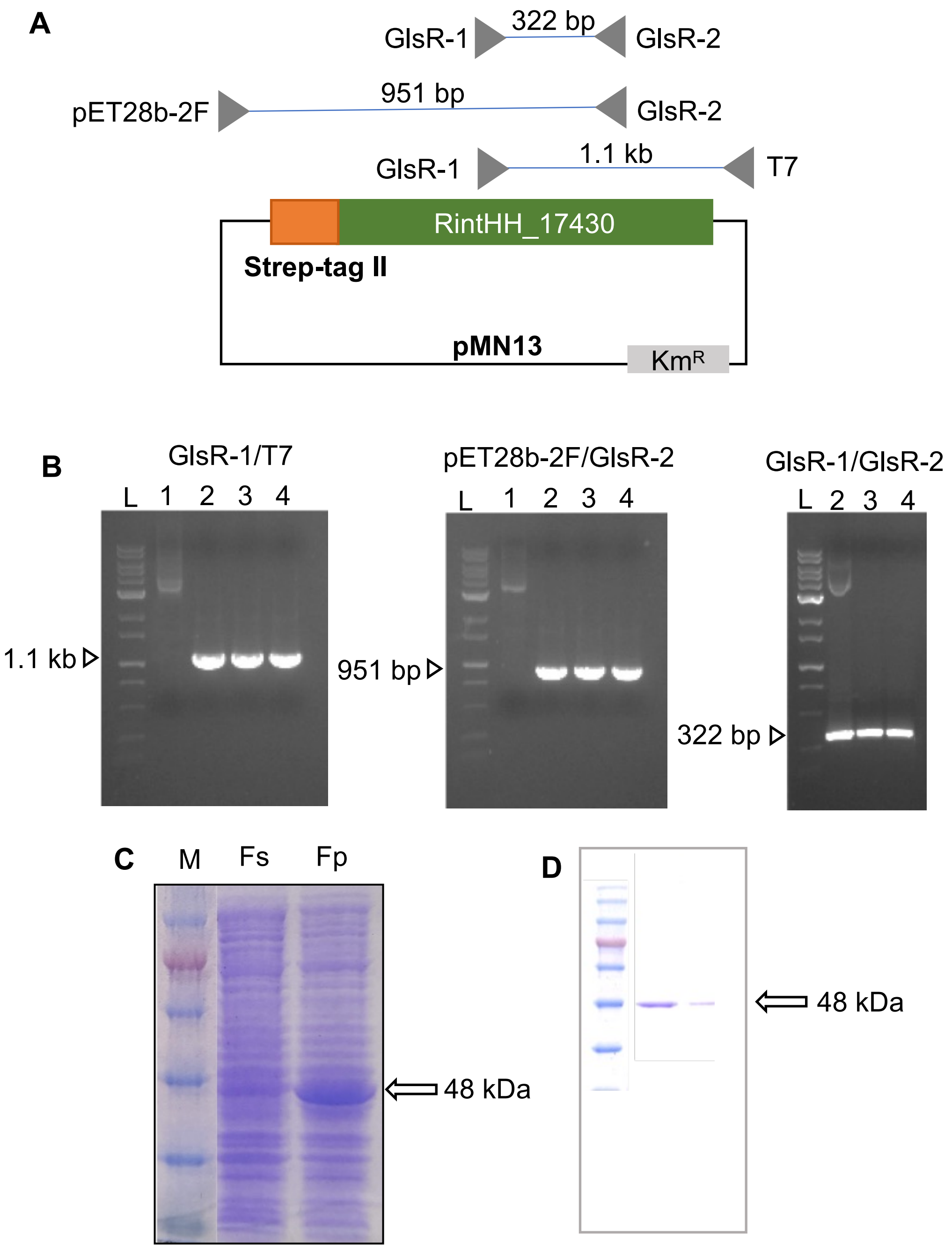


**Fig. S1. Construction of an *E. coli* strain producing RintHH_17430 from *Richelia euintracellularis*. (A)** Scheme of the *Richelia* RintHH_17430 construct cloned in pET28b(+), fused to the Strep-tag II in its N-terminus; ORF RintHH_17430 was chemically synthesized. Primers used for PCR analysis are depicted. **(B)** Verification of the construct by PCR. L, 1-kb DNA ladder. Primer pairs are indicated on top. Templates: 1, pET28b(+); 2, 3, 4, three clones of the pET28b(+)::RintHH_17430 (pMN13) construct. The plasmids were propagated in *Escherichia coli* strain BL21. **(C)** Coomassie blue-stained SDS-PAGE gels of the cell-free extract *E. coli* [pMN13] (GlsR) soluble and insoluble fraction (described in Materials and Methods). M: Size markers; Fs: Soluble fraction; Fp: Particulate fraction. **(D)** Coomassie blue-stained SDS-PAGE gel of the RintHH_17430 protein purified as described in Materials and Methods. Two different concentrations of protein were loaded.

**A**

```
>2580948285 multiple sugar transport system substrate-
binding protein [Richelia intracellularis HH01 :
CAIY01000069]
MNIQVFILRIERLSGKIKACNWQFSIVVIIYLLVFLILFGCQNLGTKNNN
QVTHVTLWQGINPPVNRDVFNKLVKKFNQTNPGIQVESIFVGEPQIPKIL
TAVVGNAPPDILLFYPQMTGKFVELGAIKSLNNWVENLPMKSEIYSNLWD
ELRLNDKIWSVPLYTSNIGIFYRPQLFSAAGIKETPETWEEFRQVAKKLT
IDRNGDGQPEQYGIVLPLGKEEWTIFCWLPFLWSAGGEIINNDNPKFDSP
EAIAALQLWQDLLKSGYAKLSAPERGYDESDFISGRVAMQITGPWTNITK
SDVDYDVFPIPASVRHATATGTGSLYVMKTTPVREKAALKFLEYILGEEF
QTEWSIKTGFIPTNEKVSKSKLYQEYASKKPGLQVFLEQMTVARARPNMA
GYSRLSDSLGRGIEAVLLGESPQKALQMAQERLRLIWNKNSK
```

**B**

**Number of amino acids:** 402

**Molecular weight:** 45355.91

**Theoretical pI:** 7.80

**Amino acid composition:**

| | | | |
|---|---|---|---|
| Ala | (A) | 25 | 6.2% |
| Arg | (R) | 15 | 3.7% |
| Asn | (N) | 25 | 6.2% |
| Asp | (D) | 15 | 3.7% |
| Cys | (C) | 2 | 0.5% |
| Gln | (Q) | 21 | 5.2% |
| Glu | (E) | 28 | 7.0% |
| Gly | (G) | 28 | 7.0% |
| His | (H) | 2 | 0.5% |
| Ile | (I) | 27 | 6.7% |
| Leu | (L) | 38 | 9.5% |
| Lys | (K) | 29 | 7.2% |
| Met | (M) | 7 | 1.7% |
| Phe | (F) | 17 | 4.2% |
| Pro | (P) | 26 | 6.5% |
| Ser | (S) | 25 | 6.2% |
| Thr | (T) | 23 | 5.7% |
| Trp | (W) | 12 | 3.0% ← |
| Tyr | (Y) | 13 | 3.2% |
| Val | (V) | 24 | 6.0% |

**Fig. S2. RintHH_17430 from *Richelia euintracellularis* HH01. (A)** Sequence of the predicted protein showing the signal peptide (lipoprotein signal peptide [Sec/SPII]; blue color) and the mature protein (green). **(B)** Amino acid composition of the mature protein showing the presence of 12 Trp residues.

**A**

```
CLUSTAL 2.1 multiple sequence alignment


RINTHH_3860     --MQPHKVILKDVEKDAWDTLEKTIIYYQGNPIGTVAALDSTANTLNYDQCFVRDFVVSA
Alr0819         MQKLNGLLTNDIIEESAWEALEKSILYYKGRPVGTVAAFDASVEALNYDQCFVRDFVSSA
Alr1521         -MKTPPINQKSLRETESWKLLESSIIYYEGNPIGTVAAQDPELAALNYDQCFLRDFVPSA
                          .  * .:*. **.:*:**:*.*:***** *.    :*******:**** **

RINTHH_3860     LLFLIKGRTEIVRNFLKQTLKLQPKKKQLGTYKPGRGLMPASFKVTFNNGQEELEADFGE
Alr0819         LIFLIKGKTDIVRNFLEETLKLQPKDRQLDAYKPGRGLIPASFKVVSDNGEEYLEADFGE
Alr1521         FVFLMDGQTDIVRNFLIETLTLQSHEKEMDCFQPGAGLMPASFKVESDGSKEYLVADFGE
                ::**:.*:*:****** :**.**.:.:::. ::** **:******  :..:* * *****

RINTHH_3860     NAIARVTPVDSSLWWIVLLRAYVVATKDYSLAYQDDFQQGIRLILDLCLATRFDMYPTLL
Alr0819         HAIARVTPVDSCLWWILLLRAYVVASKDFSLAYQPEFQTGIRLIMEICLANRFDMYPTLL
Alr1521         KAIARVPPVDSCMWWILLLRAYEKATGDLTLAREPKFQAGIKLILDLCLAHRFSMYPTML
                :*****.****.:***:*****  *: * :** : .** **:**::*** **.****:*

RINTHH_3860     VPDGACMIDRRLGIHGHPLEIQSLYYAALRAARELLICQGN-EELVFAIDNRLPVLAAHI
Alr0819         VPDGACMIDRRLGIYGHPLELQVLFYAALRAAREMLICQGN-QDVVEAIDNRLPLLCAHI
Alr1521         VPDGAFMIDRRMGVYEHPLEIQVLFYAALRAARELLLPDGDGEQYLNKVHGRLGALQYHI
                ***** *****:*:: ****:* *:*********:*: :*: :: :  :..**  *  **

RINTHH_3860     RKHYWIDLHRLNDIYRYRGEEYGKDAVNQFNIYVDSIPYSELDRWLPKGGGYLAGNVGPS
Alr0819         RQHYWIDINRLNAIYRFKSEEYGKAAVNLFNIYVDSIPYYELDKWLPKKGGYLAGNVGPS
Alr1521         RNYYWVDLKRLREIYRYKGNEFGKEIANKFNIFSQSIP-DWVIEWLPEKGGYLAGNLGPG
                *::**:*::**. ***::.*:** .* ***: :***    : .***: *******:**.

RINTHH_3860     QMDTRFFTLGNLMAVIIDLTTQEQSQAIMTLIEQRWDDLVGDMPMKITFPALENEEYKYV
Alr0819         QLDTRFFALGNLMAIISDLATEEQSQAIMTLIEDRWEDLVGDMPMKICYPALENEEYRIV
Alr1521         RMDFRFFALGNLMAILAGLASEEESQRIMNLFAHRWEDLIGYMPVKICYPALQGLEWQIV
                ::* ***:******:: .*:::*:** **.*: .**:**:* **:** :***:. *:: *

RINTHH_3860     TGCDPKNIPWSYHNAGNWPVLMWMLAAACVKTNHRELMQRALTIAQERLKNDEWPEYYDG
Alr0819         TGCDPKNIPWSYHNAGSWPVLMWMLAAASVKAGKPYIAGKAIEIAQARLLEDEWPEYYDG
Alr1521         TGCDPKNIPWSYHNGGNWPVLLWLFTAAALKTGKVELAHEAIAIAEGRLSNDKFPEYYDG
                **************.*.****:*:::**.:*:.:    :   .*: **: ** :*::******

RINTHH_3860     KQGRLIGRQSRKYQTWTIVGFLLAKEILAQPGSLSLISFDPFITKQVSQACELKFDYFDS
Alr0819         KKGRLIGKQARKYQTWTIAGFLLAAELMKNPSLLSLISFDKLPSELVSRACEFEIGSVDA
Alr1521         NNGRLIGKEARIYQTWSIAGLLVAKQFLANPDHVEFISFPDTFIGPGCSL----------
                ::*****:::* ****:*.*:*:* ::: :*. :.:***         .

RINTHH_3860     ----
Alr0819         SISR
Alr1521         ----
```

**B**

400-bp *Anabaena* P*invB*+ RintHH_3860

tttt**GGATCC**TCCTAATTCATTAAGGATATGAGGAGCCACTAAGACTTCATCTAAACCTTAAATAGCCCCAGGGATATGTT
TTTATTTGGTAAAGCATAGTTCCAGAAACAATAATGGAACGACAGATAATCTAAAAATATATCTTTATAGTACGGTTTTTA
GTAATTACTCGATAAGTTGAGGTAATTAATTGCAGTTATAGCTGTAACTTATCACAAACATCAGTGATTGCTAGTAAATTA
ATTGTGCATTCAGAATTCTAAAAATCAACGGCTTTTAAAATTTATAGTCGGCTGTTATCTCAATTAATGTAGGCTATTTAA
ACAATTATCTTTGCTATTTATGCGAAGGTAATACTACCAAAGCGCATCTACCAAAAATTCACACCCCCAATGGAGTTAATC
TGGCGatgcaaccacataaagtaattcttaaagatgtagaaaaagatgcatgggacactctagaaaaaactattatttact
atcaagggaatcctattggaacagtagcagctttagatagtacagctaatacacttaattatgaccagtgctttgtccgtg
atttgtagtttctgcattgctatttttgattaagggtagaacggaaatcgttcgcaatttcctgaaacaaaccttaaaac
tacaaccgaagaaaaacaattaggtacttataaaccaggtagaggcttaatgcccgctagttttaaagttacatttaata
atggacaagaagaattagaggctgattttggtgaaaatgccattgcgagagttacaccggttgattcttccctgtggtgga
tagttttactacgtgcttatgttgtcgcaaccaaggattattctctagcataccaagatgattttcaacaaggtattcggt
taattttggatctgtgtttagcgacacggtttgatatgtatcccaccttgctggtccctgatggagcttgcatgattgacc
gtcgcttggggattcatggtcatcctttagagattcaatctctatattatgctgctttgcgagcagcccgggaattattaa
tttgtcaggggaatgaagagttagtatttgctattgataatcgcctacccgtactggccgctcatattcgcaagcattact
ggattgatcttcatcgtttaaatgatatctatcggtatcgaggggaagaatatggtaaagatgcggtcaatcaatttaata
tatatgttgattctattccttacagcgaattggatagatggttaccaaaggggggtggttatcttgccgggaatgtggggc
catctcaaatggatacccgtttcttcactttgggtaatttaatggctgtaattattgatctaacaacacaagaacagtctc
aagcaattatgaccttgattgagcaaaggtgggatgatttagtaggagatatgccgatgaaaatcacattcccagcattgg
aaaatgaagagtataaatatgttacgggttgtgatcccaaaaatataccctggtcctatcataatgctgggaattggcctg
tcttaatgtggatgttggctgccgcctgtgttaaaaccaatcatagagaattaatgcaaagagcgcttactattgcccaag
aacgtcttaaaaatgacgagtggcctgagtattatgacggcaaacagggaagattaattggcagacaatcacggaaatatc
aaacttggacaattgtaggatttttgttggcaaaagaaatactagctcaacctggaagtctatctttgattagtttcgatc
cattcattacaaaacaggtttctcaagcttgcgagttaaaatttgactactttgactcttaa**GGATCCt**ttt

**Fig. S3.** *Richelia euintracellularis* protein RintHH_3860. **(A)** Comparison to InvB (Alr0819) and InvA (Alr1521) from *Anabaena* sp. PCC 7120. **(B)** Chemically synthesized DNA sequences: black, introduced restriction site and T tails; red, *Anabaena invB* upstream sequence; blue, RintHH_3860 sequence.

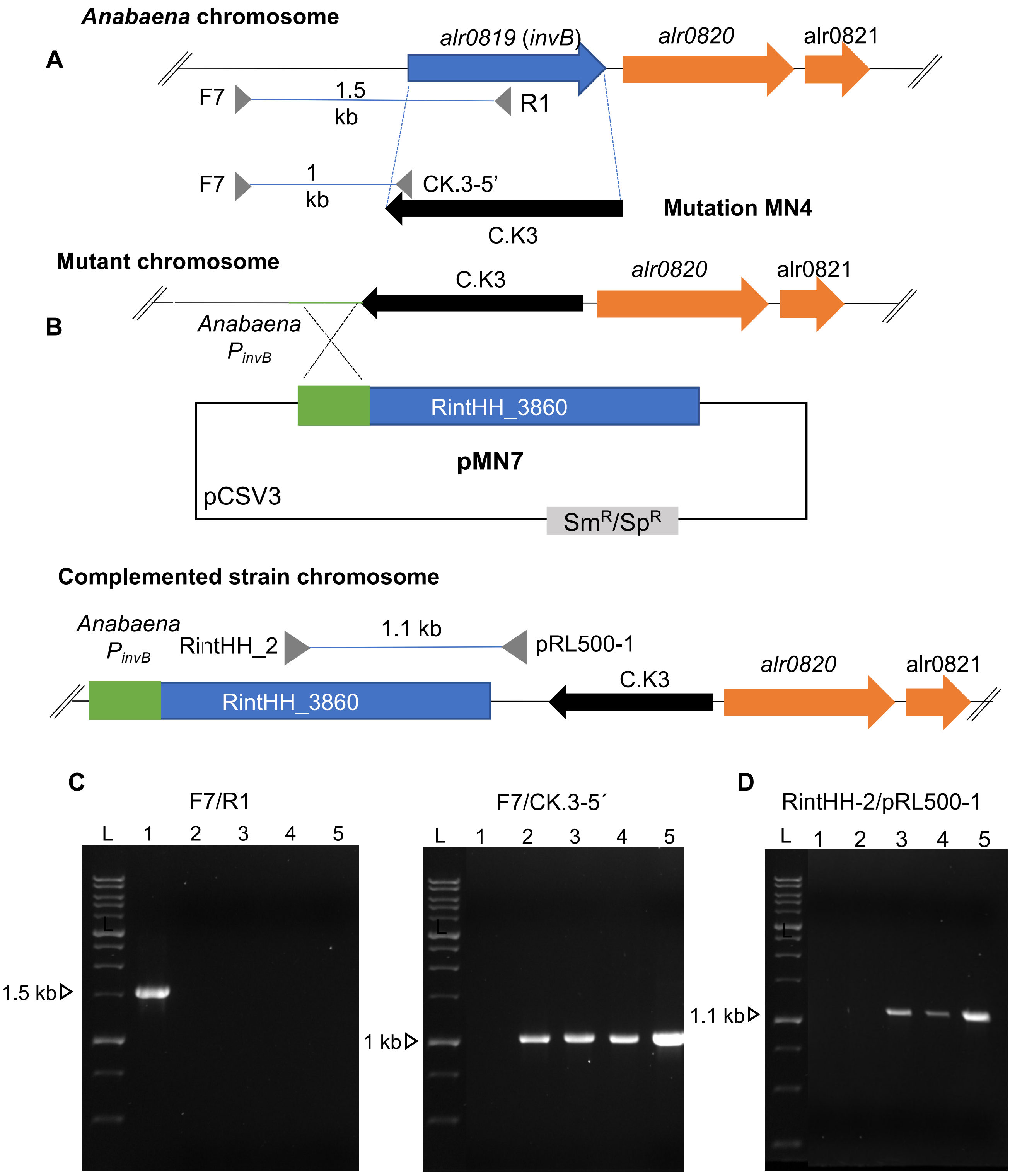


**Fig. S4. Construction and verification of an *Anabaena invB* mutant and its complementation with *Richelia euintracellularis* ORF RintHH_3860. (A)** Inactivation of *invB* in *Anabaena.* **(B)** Incorporation of the *Richelia* RintHH_3860 gene (*invB Richelia*) into the genome of the *Anabaena invB* mutant. Schematics of the genomic regions, inserted gene-cassettes and primers used for PCR are shown. **(C, D)** Verification of strains by PCR. L, 1-kb DNA ladder. Primer pairs are indicated on top. Templates: 1, wild-type *Anabaena* DNA; 2, *Anabaena invB* mutant DNA; 3, 4, 5, three independent exconjugants of the *Anabaena invB* mutant complemented with *Richelia invB*.

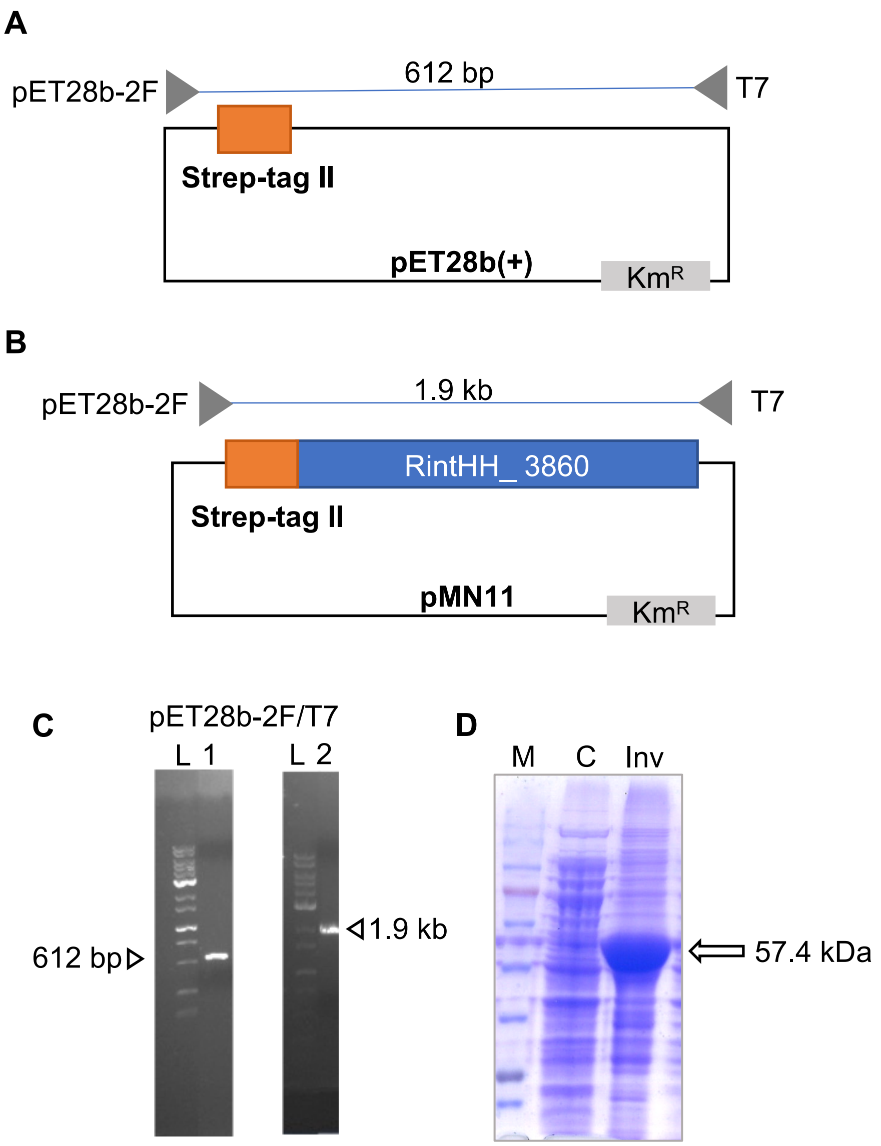


**Fig. S5. Construction of an *E. coli* strain producing RintHH_3860 from *Richelia euintracellularis.*** **(A)** Scheme of the pET28b(+) vector indicating the primers used for PCR analysis. **(B)** Scheme of the *Richelia* RintHH_3860 construct cloned in pET28b(+), fused to the Strep-tag II in its N-terminus; ORF RintHH_3860 was PCR-amplified from plasmid pMN7. Primers used for PCR analysis are depicted. **(C)** Verification of the construct by PCR. L, 1-kb DNA ladder. Primer pair is indicated on top. Templates: 1, pET28b(+) vector; 2, pET28B(+)::RintHH_3860 (pMN11). The plasmids were propagated in *Escherichia coli* strain BL21. **(D)** Coomassie blue-stained SDS-PAGE gel of the cell-free extract of an isogenic strain lacking RintHH_3860 (C: control) and cell-free extract of *E. coli* [pMN11] (Inv) as described in Materials and Methods; M, size markers.

**A** >tr|M1WZ73|M1WZ73_9NOST Branched-chain amino acid ABC transporter, amino acid-binding protein OS=Richelia intracellularis HH01 OX=1165094 GN=RINTHH_11820 PE=4 SV=1

MPRFCTAMFTSITVLVVSLLTVACVPQSTTNSNTNTRTTNTKSKGLKIGSLLPTTGDLAS
VGQQMAGAVTLLVDTINDCGGVNGEQVSLVEVDSQTDPRAGAAGMTKLATLDKVGGVVGA
FASSVSSAAVSIAVPNKVMLVSPGSTSPIFTDNSQKGYYKGFWARTAPPDTYQALALAQL
ARKKGFTRVSTAVINNDYGVGFEKAFVQAFEKLGGTVVNKYKPVRYDPKAQTFDTEAASV
FASSPEAVIAVLYAETGSLFLKAAYQQGLTEGVQIMLTDGVKSDSFPKQVGQTGDGKYII
SEAVGTVPGSNGKALDALNKLWREKKGNPPGEYAPQVWDAVALLTLAAQAAKDNSGLGIS
NKIKEVANPPGKEVTDVCQGLKLLKEGKDINYQGASGNVDIDENGDVIGVYNVWTVDSNG
KIQVIDKVSPNTTVKMRS

Trp residues in mature protein: 4

**B** >tr|M1WZB0|M1WZB0_9NOST Extracellular solute-binding protein, family 3 OS=Richelia intracellularis HH01 OX=1165094 GN=RINTHH_12770 PE=3 SV=1

MFMFKSALSSVITVLTISLIACSSNTVTPGKNNSLVNRIKNRGRVICGVSGEIPGFSFVD
TDGRYRGLDVDVCRAMAAALFDKPDAVDYRNLNAKERFTAVQTGEVDLLSRNTTLTISRD
TSVGMAFGPIVFYDGQGIMVSKRSKVKSLKDLNGKAICTQTGTTNEQNLADKMKLLGINY
KPVVFEDINTAFATYQQGRCSAITSDKSQLISRRTTLPERENHIILGESLSQEPLAPAVA
DGDAKWADALKWVIYALIKAEELGITSQNVMQKINSDNPEINRLLGNGSNLGEGMGLSND
FVVRVIKHVGNYGEIYDRNLGLKTELNLPRSYNRQWMKGGLLYAPPFR

Trp residues in mature protein: 3

**C** >tr|M1WZJ9|M1WZJ9_9NOST ABC transporter, periplasmic spermidine putrescine-binding protein PotD (TC 3.A.1.11.1) OS=Richelia intracellularis HH01 OX=1165094 GN=RINTHH_7180 PE=4 SV=1

MQSVLTLATLQLAVGCSSKQKTQFTVNLLKNSIPAHLINKFSQSLKQNIQLEFSPVEQLQ
TIFQELIDWQQLSKNNTRKGLNNPPLFWPSKSNKVADVVTLGNYWLDIAIKQGLIQPLDR
EQIPNWHHLPKEWQRLVMRNQQGDLDRQGQIWAAPYRWGTTMIVYRRDKFDRLGWEPQDW
GDLWREELRDRISILDHPREVIGLVLKRLGKSYNYEDLEFITELEPLLHSLHQQVKFYSS
DKYLEALLMKDTWLAVGWSSDILRAIARNTTLAAVIPKSGTALWADMWACPYTGVSANIN
KKAIHQWINFCWQAENAKQIALLTKTNSPIPNYIKSDEIQKSLREISLNDQIFQKSEFIH
CLEPTLNAKYESLFTRITKA

Trp residues in mature protein: 16

**Fig. S6. RintHH_11820 (A), RintHH_12770 (B), RintHH_7180 (C) from *Richelia euintracellularis* HH01.** Shown is the sequence of each predicted protein indicating the signal peptide (lipoprotein signal peptide [Sec/SPII]; green color) and the mature protein (blue). The number of Trp residues in each mature protein is also indicated.

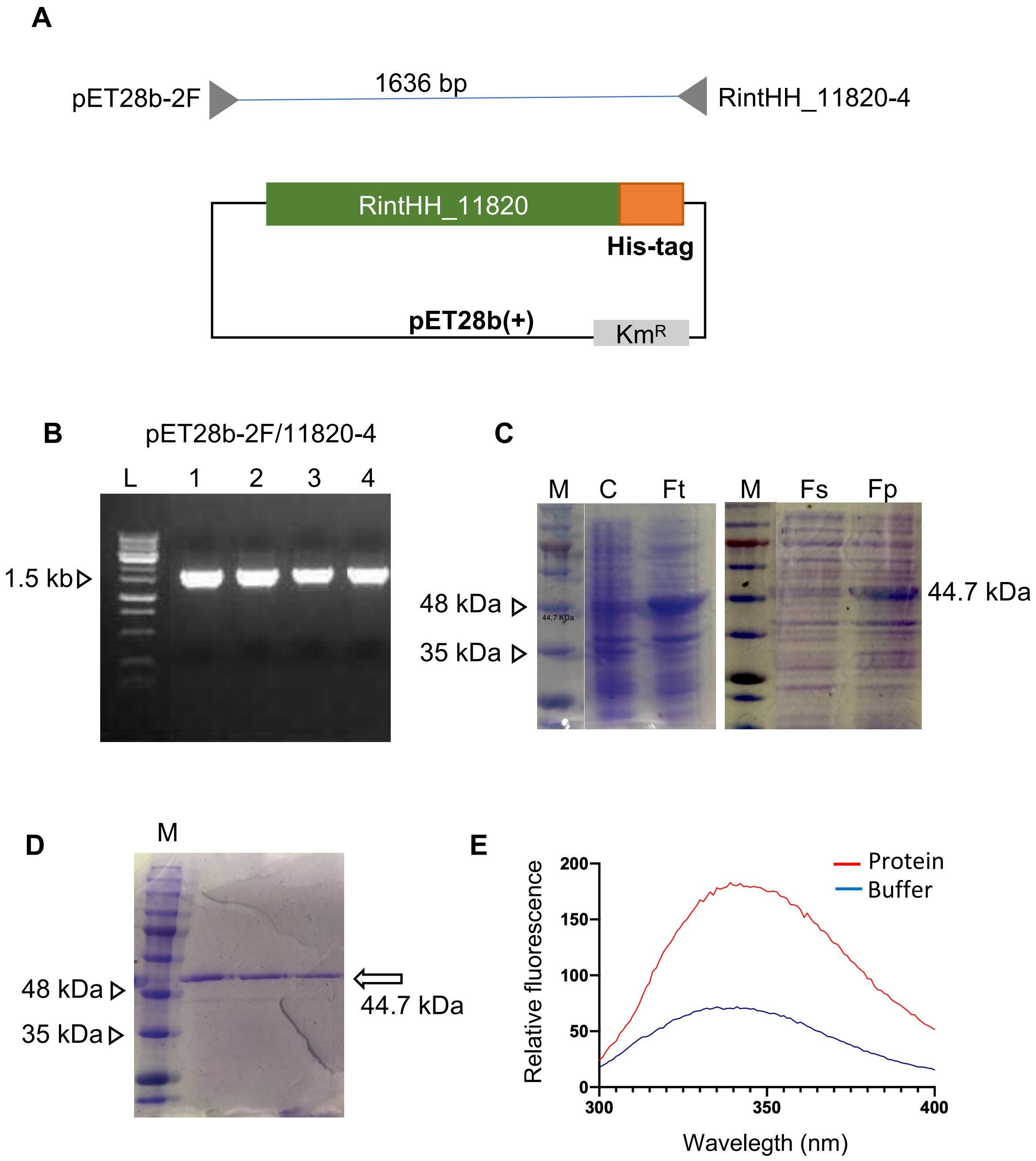


**Fig. S7. Construction of an *E. coli* strain producing RintHH_11820 from *Richelia euintracellularis*. (A)** Scheme of the *Richelia* RintHH_11820 construct cloned in pET28b(+), fused to a 6xHis tag in its C-terminus; ORF RintHH_11820 was chemically synthesized. Primers used for PCR analysis are depicted. **(B)** Verification of the construct by PCR. L, 1 kb DNA ladder; 1, 2, 3, 4, four clones of the pET28b(+)::RintHH_11820 construct. The plasmids were propagated in *E. coli* strain BL21. **(C)** Coomassie blue-stained SDS-PAGE gels showing extracts of clone 2 [pET28b(+)::RintHH_11820]. M, size markers; C, total fractions of non-induced culture, C; Ft, cell free-extract of induced culture; Fs, soluble fraction; Fp, particulate fraction. **(D)** Coomassie blue-stained SDS-PAGE gel of the RintHH_11820 protein purified as described in Materials and Methods. Three different concentrations of protein were loaded. Note that RintHH_11820 shows altered motility. **(E)**, Intrinsic tryptophan fluorescence(excitation, 280 nm) of RintHH_11820 protein purified in comparison with the buffer signal.

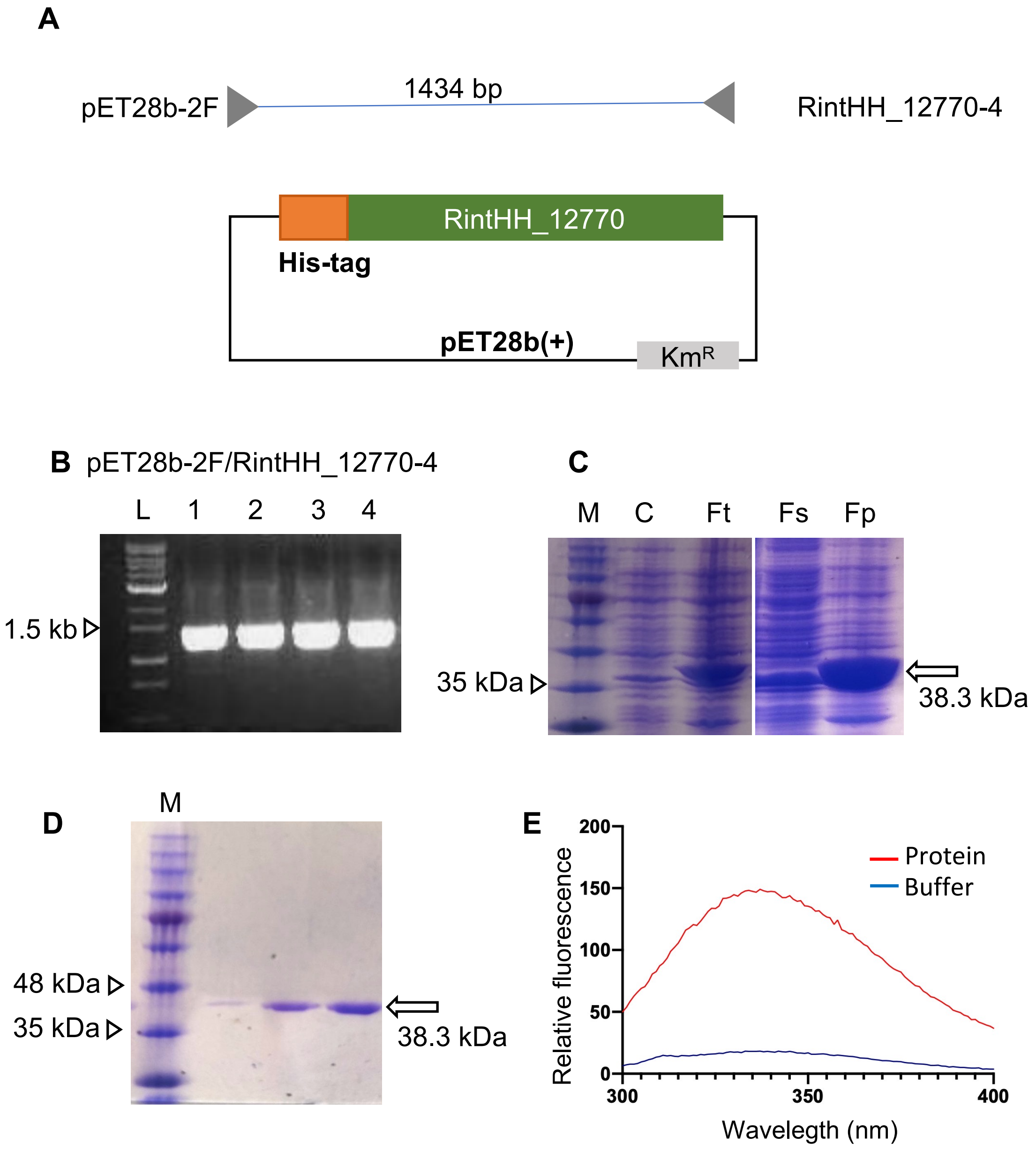


**Fig. S8. Construction of an *E. coli* strain producing RintHH_12770 from *Richelia euintracellularis*. (A)** Scheme of the *Richelia* RintHH_12770 construct cloned in pET28b(+), fused to a 6xHis tag in its N-terminus; ORF RintHH_12770 was chemically synthesized. Primers used for PCR analysis are depicted. **(B)** Verification of the construct by PCR. L, 1 kb DNA ladder; 1, 2, 3, 4, four clones of the pET28b(+)::RintHH_12770 construct. The plasmids were propagated in *E. coli* strain BL21. **(C)** Coomassie blue-stained SDS-PAGE gels showing extracts of clone 2 [pET28b(+)::RintHH_12770]. M, size markers; C, total fractions of non induced cultures; Ft, cell-free extract of induced culture; Fs, soluble fraction; Fp, particulate fraction. **(D)** Coomassie blue-stained SDS-PAGE gel of the RintHH_12770 protein purified as described in Materials and Methods. Three different concentrations of protein were loaded. **(E)** Intrinsic tryptophan fluorescence (excitation, 280 nm) of purified RintHH_12770 protein in comparison with the buffer signal.

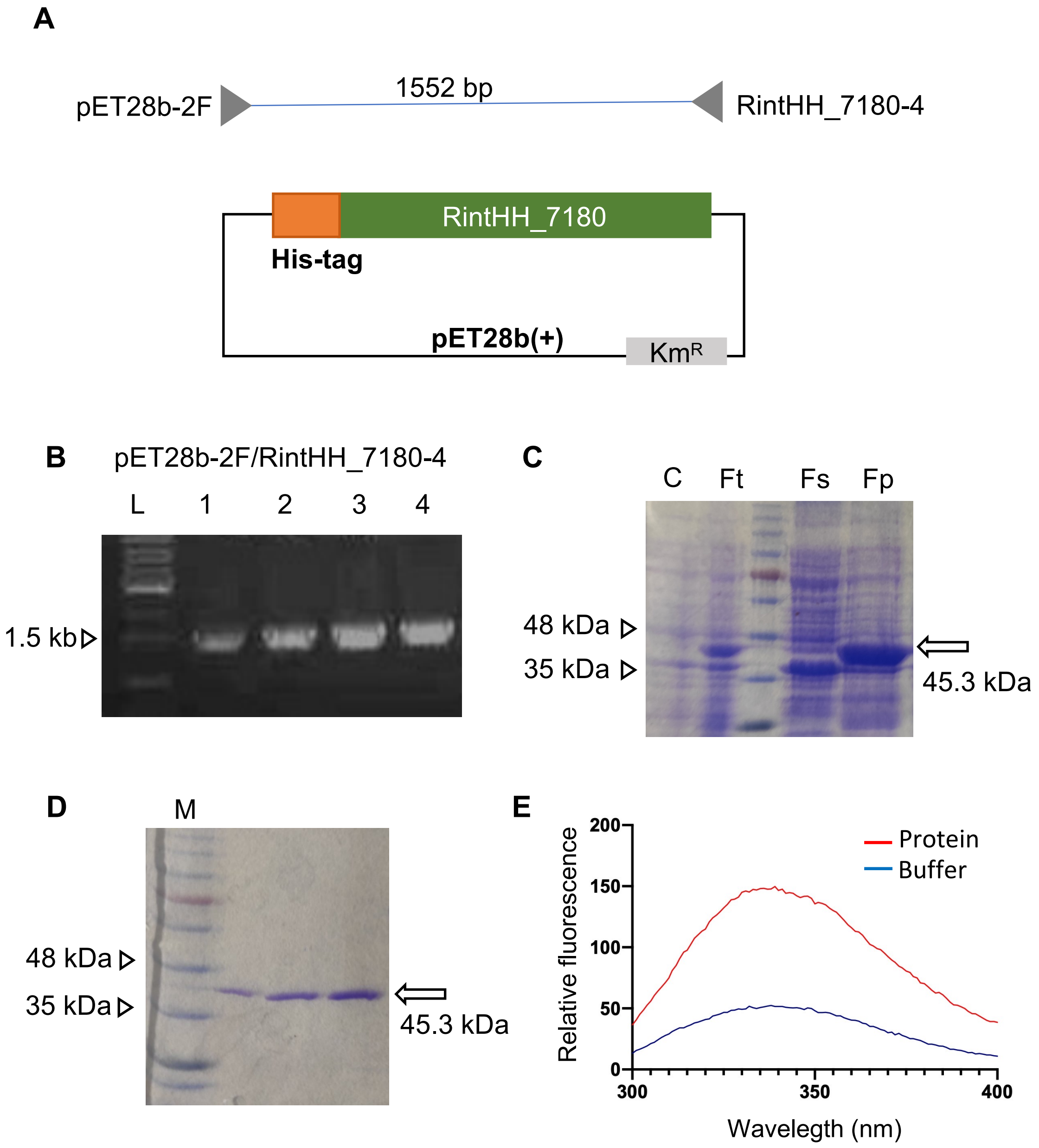


**Fig. S9. Construction of an *E. coli* strain producing RintHH_7180 from *Richelia euintracellularis*. (A)** Scheme of the *Richelia* RintHH_7180 construct cloned in pET28b(+), fused to a 6xHis tag in its N-terminus; ORF RintHH_7180 was chemically synthesized. Primers used for PCR analysis are depicted. **(B)** Verification of the construct by PCR. L, 1 kb DNA ladder; 1, 2, 3, 4, four clones of the pET28b(+)::RintHH_7180 construct. The plasmids were propagated in *E. coli* strain BL21. **(C)** Coomassie blue-stained SDS-PAGE gels showing extracts of clone 2 [pET28b(+)::RintHH_7180]. C, total fractions of non-induced culture; Ft, cell-free extract of induced culture; Fs, soluble fraction; Fp, particulate fraction. **(D)** Coomassie blue-stained SDS-PAGE gel of the RintHH_7180 protein purified as described in Materials and Methods. Three different concentrations of protein were loaded; M, size markers. **(E)** Intrinsic tryptophan fluorescence (excitation, 280 nm) of purified RintHH_7180 protein in comparison with the buffer signal.

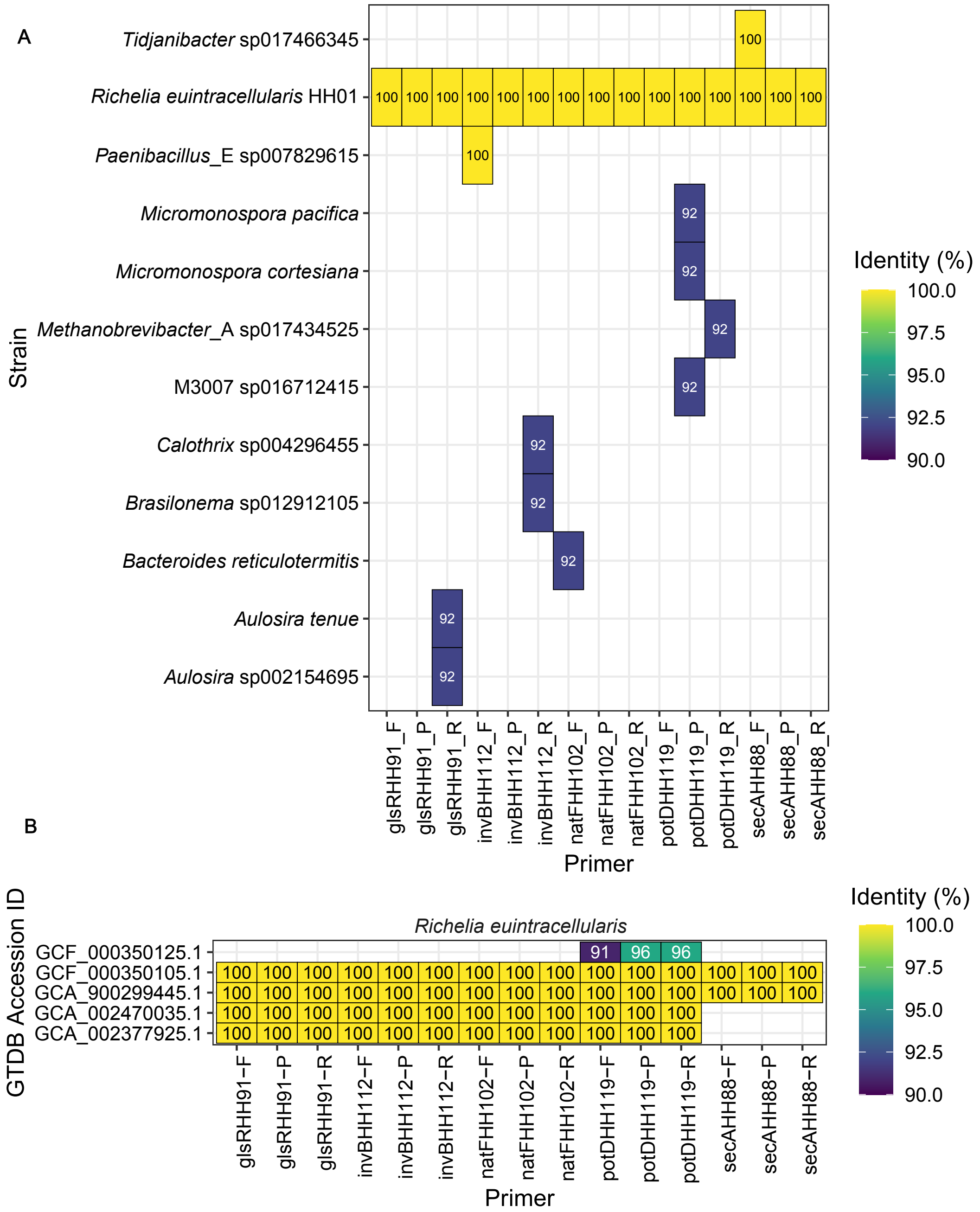


**Fig. S10. Results from BLASTn analyses for testing specificity of RT-qPCR assays against *Richelia euintracellularis* HH01. (A)** The BLASTn results using Database I (Representative genomes) indicate the RT-qPCR assays are highly specific against *Richelia euintracellularis* HH01. **(B)** BLASTn results using Database II (Cyanobacteria). The GTDB accession ID indicates which genome the aligned sequence belongs to while the primer axis indicates which of our designed primers was used as query. The results have been filtered and selected according to the criteria mentioned in the Suppl. methods. The "GCF_000350125.1" represents *R. euintracellularis* strain HM01 while the rest of the IDs belong to strain *R. euintracellularis* HH01. The primer names are in the structure of 'gene name', 'length of amplified sequence' and forward (F), reverse (R) or probe (P).

### potD primers

potD forward primer alignment for (*R. intracellularis* HM01 = GCF_000350125.1 in Figure S10B)

```
>FDBDIGKM_00912 hypothetical protein [GCF_000350125.1
d__Bacteria;p__Cyanobacteria;c__Cyanobacteriia;o__Cyanobacteriales;f__Nostocaceae;g__Richelia;s
__Richelia
intracellularis]
Length=1164

 Score = 31.9 bits (34),  Expect = 6.0
 Identities = 20/22 (91%), Gaps = 0/22 (0%)
 Strand=Plus/Plus

Query  1    CGCAACCAGCAAGGTGATTTAG  22
            |||||||||||||||  | ||||
Sbjct  433  CGCAACCAGCAAGGTAAGTTAG  454
```

potD probe alignment for (*R. intracellularis* HM01 = GCF_000350125.1 in Figure S10B)

```
>FDBDIGKM_00912 hypothetical protein [GCF_000350125.1
d__Bacteria;p__Cyanobacteria;c__Cyanobacteriia;o__Cyanobacteriales;f__Nostocaceae;g__Richelia;s
__Richelia
intracellularis]
Length=1164

 Score = 41.9 bits (45),  Expect = 0.012
 Identities = 24/25 (96%), Gaps = 0/25 (0%)
 Strand=Plus/Plus

Query  1    TCGTCAGGGCCAGATTTGGGCAGCA  25
            || ||||||||||||||||||||||
Sbjct  456  TCATCAGGGCCAGATTTGGGCAGCA  480
```

potD reverse primer alignment for (*R. intracellularis* HM01 = GCF_000350125.1 in Figure S10B)

```
>FDBDIGKM_00912 hypothetical protein [GCF_000350125.1
d__Bacteria;p__Cyanobacteria;c__Cyanobacteriia;o__Cyanobacteriales;f__Nostocaceae;g__Richelia;s
__Richelia
intracellularis]
Length=1164

 Score = 41.9 bits (45),  Expect = 0.012
 Identities = 24/25 (96%), Gaps = 0/25 (0%)
 Strand=Plus/Minus

Query  1    TGTGGTTCCCAACCTAATCTATCAA  25
            ||||||| |||||||||||||||||
Sbjct  551  TGTGGTTTCCAACCTAATCTATCAA  527
```

**Fig. S11. Results of BLASTn analyses between oligonucleotides deisgned to detect ReuHH01 *potD* as queries with the *Richelia euintracelluarlis* HM01 genome**. Alignments are shown for the forward (top), probe (middle), and reverse primer (bottom). Sequence similarities are shown in Fig. S10B.
